# DNA Methylation Signature on Phosphatidylethanol, not Self-Reported Alcohol Consumption, Predicts Hazardous Alcohol Consumption in Two Distinct Populations

**DOI:** 10.1101/820910

**Authors:** Xiaoyu Liang, Amy C. Justice, Kaku So-Armah, John H. Krystal, Rajita Sinha, Ke Xu

## Abstract

The process of diagnosing hazardous alcohol drinking (HAD) is based on self-reported data and is thereby vulnerable to bias. There has been an interest in developing epigenetic biomarkers for HAD that might complement clinical assessment. Because alcohol consumption has been previously linked to DNA methylation (DNAm), here, we aimed to select DNAm signatures in blood to predict HAD from two demographically and clinically distinct populations (N_total_=1,549). We first separately conducted an epigenome-wide association study (EWAS) for phosphatidylethanol (PEth), an objective measure of alcohol consumption, and for self-reported alcohol consumption in Cohort 1. We identified 102 PEth-associated CpGs, including 32 CpGs previously associated with alcohol consumption or alcohol use disorders. In contrast, no CpG reached epigenome-wide significance on self-reported alcohol consumption. Using a machine learning approach, two subsets of CpGs from EWAS on PEth and on self-reported alcohol consumption from Cohort 1 were separately tested for the prediction of HAD in Cohort 2. We found that a subset of 130 CpGs selected from the EWAS on PEth showed an excellent prediction of HAD with area under the ROC curve (AUC) of 91.31% in training set and 70.65% in validation set of Cohort 2. However, CpGs preselected from the EWAS on self-reported alcohol consumption showed a poor prediction of HAD with AUC 75.18% in the training set and 57.60% in the validation set. Our results demonstrate that an objective measure for alcohol consumption is a more informative phenotype than self-reported data for revealing epigenetic mechanism. The PEth-associated DNAm signature in blood is a robust biomarker for alcohol consumption.

## INTRODUCTION

Hazardous alcohol drinking (HAD) is detrimental to health and is highly correlated with medical comorbidities and psychiatric diseases ^1, 2^. Diagnosing HAD is challenging due to a lack of stable and objective measures for chronic heavy alcohol consumption ^3^. Phosphatidylethanol (PEth) is a lipid metabolite of ethanol formed from phosphatidylcholine in erythrocytes and has been proposed as a biomarker for alcohol consumption. Compared with self-reported data, PEth reliably detects ethanol levels up to 21 days after the last drink ^4^, and the PEth level is highly correlated with alcohol consumption ^5^. However, the clinical applicability of PEth is limited because its half-life is approximately 4–7 days ^6^. Thus, other more stable biomarkers for alcohol consumption are needed to inform clinical practice.

Epigenetic signatures have emerged as attractive biomarkers for complex diseases such as cancers and neurodegenerative diseases ^7^. Epigenetic markers may reflect environmental exposures, including alcohol consumption. Among these epigenetic markers, DNA methylation (DNAm) biomarkers are particularly attractive because they are relatively stable and capture an early stage of pathophysiological changes ^8, 9^. A recent longitudinal study on DNAm showed that most DNA methylome changes occurred 80-90 days before clinically detectable glucose elevation ^10^, suggesting that DNAm is involved in an early stage of diabetes. Finally, epigenetic modifications can be reliably detected in noninvasive fluids and biospecimens ^11^. Thus, the utility of epigenetic alterations has motivated the biomarker research field to develop epigenetic signatures derived from easily accessible cells for clinical use ^12–14^.

DNAm markers are emerging as diagnostic biomarkers in many areas of medicine and are applied to predict complex diseases ^15^. For example, DNAm markers on the promoters of several genes, including *BMP3, NDRG4,* and *SPEPT9,* in blood or stool samples have been approved by the Food and Drug Administration as biomarkers for colorectal cancer screening ^16^. DNAm markers on *APP*, *BACE1*, and *LEP1* in erythrocytes have been applied in predicting the prognosis of Alzheimer’s disease ^17^. DNAm markers also distinguish smokers and nonsmokers ^18, 19^. However, we do not yet have validated DNAm biomarkers for the diagnosis of HAD.

Recent studies have shown that alcohol consumption modifies DNAm ^20^ in animals and in the human epigenome from blood, liver, and saliva cells ^18, 21–25^. Epigenome-wide association studies (EWAS) have identified hundreds of DNAm cytosine-phosphate-guanine sites (CpGs) from blood samples that are associated with alcohol consumption ^26–29^, alcohol use disorders ^30, 31^, stress-related alcohol consumption ^32^, and fetal alcohol syndrome ^33–36^. A large number of CpGs in the human leukocyte DNA methylome have recently been reported to have associations with dietary folate and alcohol intake ^37^. Some CpGs have been found to be associated with alcohol consumption in different cell types, ethnic groups, and phenotypic assessments ^29, 30, 38^. Among the reported CpGs for alcohol consumption, more than a dozen CpGs have been replicated. For example, cg11376147 on *SLC43A1* has been linked to alcohol consumption and HAD diagnosis in several studies ^18, 29, 30^. Thus, DNAm in blood has been proposed as a diagnostic and prognostic biomarker of alcohol consumption for clinic use ^39^. For this purpose, a previous study identified a panel of 114 CpGs as biomarkers for alcohol consumption ^30^. However, these CpGs have not been validated in independent studies.

One of the limitations of previous EWAS is that alcohol consumption was assessed by self-report, which may lead to inaccurate assessment and introduce bias ^30, 40, 41^. A self-reported phenotype may, in part, explain the discrepancy of EWAS findings on alcohol consumption or alcohol use-related phenotypes observed in previous studies. Objective measures such as PEth may improve the association signals for alcohol consumption in EWAS because PEth-associated DNAm markers are more proximal to the biological changes and pathological processes underlying HAD.

In this study, we hypothesized that the DNAm signatures associated with PEth would be a more robust predictor of HAD than self-reported drinking data. We conducted a 2-stage study with the goal of identifying PEth-associated DNAm CpGs and then linking the PEth-associated methylation features to HAD (N_total_ = 1,549). We compared the findings of DNAm markers for PEth with those for self-reported alcohol consumption. The first stage included an EWAS for PEth in a discovery sample and in a replication sample from Cohort 1. An EWAS of self-reported Alcohol Use Disorders Identification Test-Consumption (AUDIT-C, first 3-items of AUDIT) score in the same individuals of Cohort 1 were also conducted in comparison of the EWAS findings on PEth. In the second stage, we applied a recently developed machine learning method, elastic net regularization (ENR), to select CpGs for predicting HAD defined by a self-reported 10-item AUDIT measurement in a demographic and clinically independent sample (Cohort 2). The preselected PEth-associated CpGs from the EWAS of PEth in Cohort 1 were optimized to predict HAD in Cohort 2. Using the same analytic approach, the HAD predicting procedure using the preselected CpGs from the EWAS on AUDIT-C score was also performed. The analytical strategy is presented in **Figure 1**.

**Figure 1.**
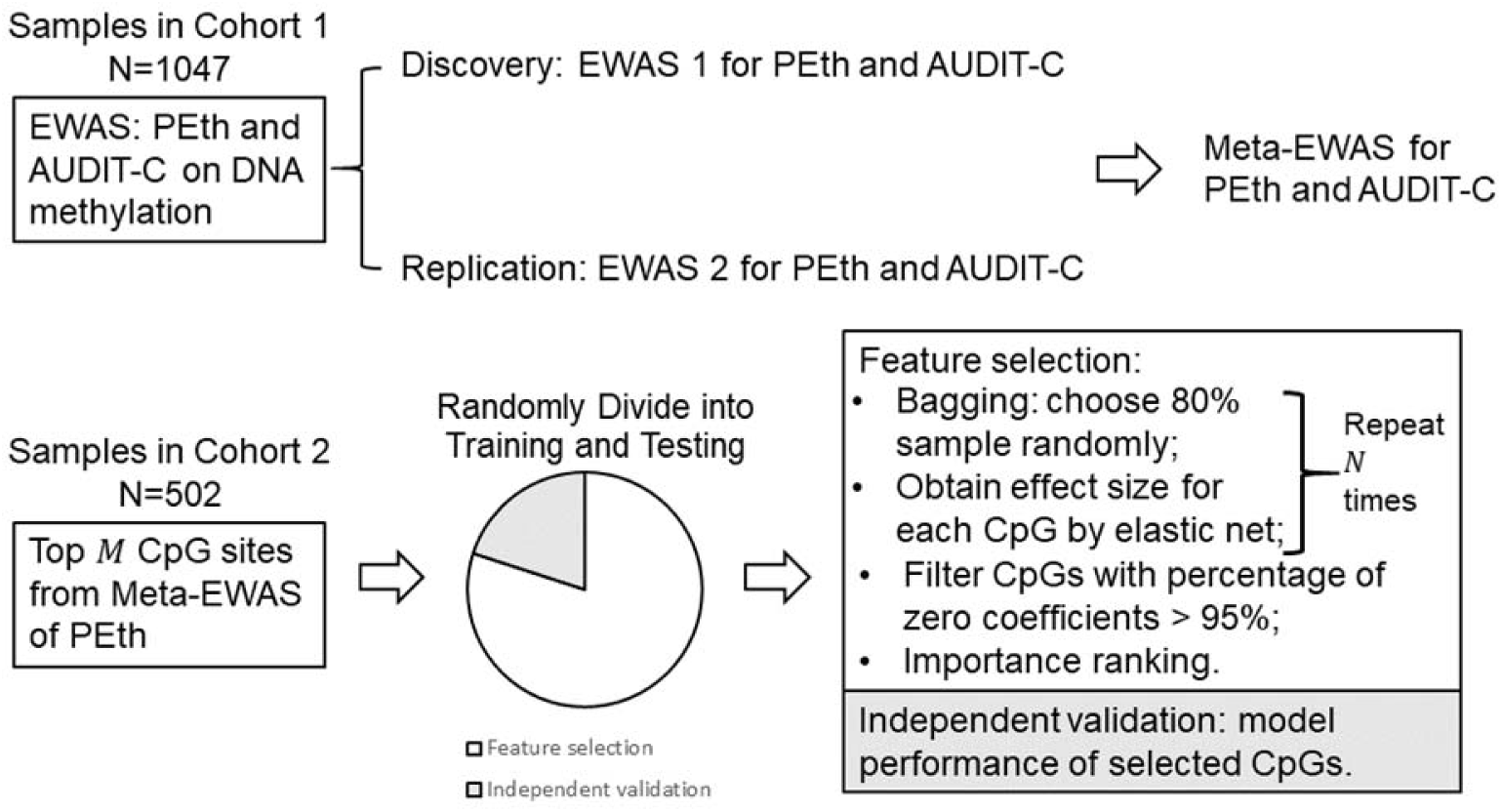
Study design for the epigenome-wide association study for alcohol consumption.

## MATERIALS AND METHODS

### Sample descriptions

**Cohort 1 (N=1,047):** The DNA samples in Cohort 1 were from the Veterans Aging Cohort Study (VACS). The VACS is a longitudinal cohort of HIV-positive and HIV-negative participants seen in infectious disease and general medical clinics. The study is funded primarily by the National Institute on Alcohol Abuse and Alcoholism at the National Institutes of Health ^42^. Data were obtained from the patients after they provided written consent; data were collected via telephone interviews, focus groups, and full access to the national Veterans Affairs (VA) electronic medical record system. All subjects in this subset of the VACS cohort were men.

Samples in Cohort 1 were divided into a discovery set (N=580) and a replication set (N=467) for EWAS and for predicting HAD. A majority of discovery samples were HIV-positive (∼85.34%), and all replication samples were HIV-positive. HIV Viral Load (VL) was measured per standard of care by polymerase chain reaction as copies per milliliter. The adherence to antiretroviral therapy (ART adherence) was obtained by a survey in the same time window as the blood draw for the measurement of DNA methylation. Genomic DNA was extracted from whole blood using a standard method ^12^.

**Cohort 2 (N=502)**: We recruited 502 HIV-negative healthy community volunteers who responded to advertisements placed either online or in local newspapers and at a community center in New Haven, CT ^43^. The subjects were 18–50 years old. We excluded subjects who met the Diagnostic and Statistical Manual of Mental Disorders, 4th Edition (DSM-IVTR) (American Psychiatric Association, 1994) criteria for substance dependence on any drug or alcohol other than nicotine. Subjects with head injury or those who used prescribed medications for any psychiatric or medical disorders were also excluded. Women on oral contraceptives, women who were peri- and postmenopausal, women who had a prior hysterectomy and women who were pregnant, or lactating were excluded. Participants also received a physical examination during a separate session by a research nurse who assessed cardiovascular, renal, hepatic, pancreatic, hematopoietic, and thyroid functions to ensure that all participants were in good health. A breathalyzer test and urine toxicology screens were conducted at each appointment to ensure a drug-free status among participants. Cohort 2 was divided into a training set and a testing set for machine learning prediction of HAD. Genomic DNA was extracted from whole blood using a standard method.

All phenotypic data in Cohort 1 and Cohort 2 were obtained in the same time window as the blood draws for DNA methylation profiling. The study was approved by the committee of the Human Research Subject Protection at Yale University and the IRB committee of the Connecticut Veteran Healthcare System.

### Phosphatidylethanol (PEth) measurement

In this study, PEth was only measured in Cohort 1 using dried blood spot samples derived from frozen peripheral blood mononuclear cells stored at -80°C ^5^. PEth can be detected at concentrations as low as 2 ng/ml. A study showed that the PEth value is linearly related to alcohol consumption ^44^. In forensics, 20 ng/ml of PEth was used as a cutoff to detect harmful alcohol use ^45^. The sensitivity of PEth has been reported to be 99% ^44^, with several studies showing the assay to have perfect specificity, including in the presence of liver disease and hypertension. We previously reported that PEth was highly correlated with the AUDIT-C score from electronic records ^46^.

### Definition of hazardous alcohol drinking (HAD)

In Cohort 1, HAD was defined by a PEth level greater than 20 ng/ml and an AUDIT-C score greater than 4. In Cohort 2, HAD was defined by a 10-item AUDIT score greater than 8 for men and greater than 7 for women. Demographic and clinical variables for HAD versus non-HAD participants in Cohort 1 and Cohort 2 are presented in **Table 1**.

**Table 1.** Demographic and clinical characteristics for Cohort 1 and Cohort 2

|  | Cohort 1: |  | Cohort 1: |  | Cohort 2 |  |
| --- | --- | --- | --- | --- | --- | --- |
|  | Discovery Sample |  | Replication Sample |  |  |  |
|  | HAD | non-HAD | HAD | non-HAD | HAD | non-HAD |
| | PEth ≥ 20<br>(N = 166) | PEth $<$ 20<br>(N = 414) | PEth ≥ 20<br>(N = 135) | PEth $<$ 20<br>(N = 332) | Men:<br>AUDIT ≥ 8<br>Women:<br>AUDIT ≥ 7<br>(N = 150) | Men:<br>AUDIT $<$ 8<br>Women:<br>AUDIT $<$ 7<br>(N = 333) |
| Age (year) | 49.28 ± 7.25 | 49.25 ± 8.13 | 47.50 ± 7.08 | 48.18 ± 8.03 | 26.77 ± 7.08 | 29.61 ± 9.38<sup>a</sup> |
| Sex (male, %) | 100 | 100 | 100 | 100 | 64.67 | 35.74<sup>b</sup> |
| Race (AA, %) | 90.36 | 79.71<sup>c</sup> | 82.22 | 81.02 | 12.08 | 22.22<sup>c</sup> |
| Smoker (%) | 70.91 | 53.92<sup>d</sup> | 63.64 | 54.91 | 39.33 | 13.21<sup>d</sup> |
| Alcohol (AUDIT-C) | 4.73 ± 2.65 | 2.57 ± 2.40<sup>e</sup> | 4.80 ± 2.30 | 2.28 ± 2.24<sup>e</sup> | NA | NA |
| HIV-infection (%) | 88.55 | 84.54 | 100 | 100 | NA | NA |
| VL (log10) | 2.85 ± 1.24 | 2.6 ± 1.2 | 2.69 ± 1.20 | 2.68 ± 1.24 | NA | NA |
| ART adherence (%) | 69.23 | 81.69<sup>f</sup> | 72.73 | 77.2 | NA | NA |
| CD4+ T (%) | 0.06 ± 0.06 | 0.07 ± 0.06 | 0.10 ± 0.05 | 0.09 ± 0.04 | 0.18 ± 0.05 | 0.18 ± 0.05 |
| CD8+ T (%) | 0.17 ± 0.09 | 0.16 ± 0.09 | 0.18 ± 0.09 | 0.18 ± 0.08 | 0.10 ± 0.04 | 0.09 ± 0.04 |
| NK (%)<sup>h</sup> | 0.07 ± 0.05 | 0.08 ± 0.06 | 0.09 ± 0.03 | 0.08 ± 0.03 | 0.03 ± 0.03 | 0.03 ± 0.03 |
| B cell (%)<sup>h</sup> | 0.08 ± 0.05 | 0.09 ± 0.05<sup>g</sup> | 0.08 ± 0.03 | 0.08 ± 0.04 | 0.07 ± 0.03 | 0.07 ± 0.03 |
| Monocyte (%)<sup>h</sup> | 0.12 ± 0.04 | 0.11 ± 0.04 | 0.11 ± 0.04 | 0.11 ± 0.03 | 0.08 ± 0.02 | 0.08 ± 0.02 |
| Granulocyte (%)<sup>h</sup> | 0.53 ± 0.12 | 0.53 ± 0.14 | 0.50 ± 0.11 | 0.50 ± 0.12 | 0.58 ± 0.09 | 0.59 ± 0.09 |

AA: African American, AUDIT: Alcohol Use Disorders Identification Test, AUDIT-C: first three questions of the Alcohol Use Disorders Identification Test, VL: viral load, ART: antiretroviral therapy
<sup>a</sup>two-sample t-test P-value = 2.83E-04
<sup>b</sup>chi-square test P-value = 5.94E-09
<sup>c</sup>chi-square test P-value < 1.27E-02
<sup>d</sup>chi-square test P-value < 2.65E-04
<sup>e</sup>two-sample t-test P-value < 3.50E-14
<sup>f</sup>chi-square test P-value = 3.69E-03
<sup>g</sup>two-sample t-test P-value = 2.34E-02
<sup>h</sup>Cell type compositions estimated by methylation

### DNA methylation and data quality control

In Cohort 1, DNAm for the discovery sample was profiled by using the Illumina Infinium HumanMethylation450 Beadchip (Illumina HM450K) (San Diego, CA, USA). DNAm for the replication sample was assessed by using the Illumina Infinium MethylationEPIC Beadchip (Illumina EPIC) (San Diego, CA, USA). In Cohort 2, DNAm was measured by using Illumina HM450K. All samples in Cohorts 1 and 2 were processed at the Yale Center for Genomic Analysis^12^.

Methylation raw data were retrieved using the minfi R package (version 1.18.1), and downstream analyses were performed using minfi and R. We performed the probe normalization and batch-correction procedure using the pipeline reported by Lehne *et al.*^47^. We removed CpGs on sex chromosomes and CpGs within 10 base pairs of single nucleotide polymorphisms. In Cohort 1, only common CpGs between the Illumina HM450K and Illumina EPIC array were analyzed in meta-analyses. After QC, a total of 408,583 CpGs remained for analysis. We also compared the predicted sex with self-reported sex. All samples were matched as male. In Cohort 2, we applied the same QC criteria. A total of 437,722 CpGs remained for analysis. Methylation inferred sex matched with self-reported sex data in this cohort.

Six cell types (CD4+ T cells, CD8+ T cells, NK T cells, B cells, monocytes, and granulocytes) in the blood were estimated in each sample for both cohorts using the method described by Houseman *et al*. ^48, 49^.

### Discovery and replication EWAS in Cohort 1

EWAS were separately performed to test the association of each CpG methylation with PEth and AUDIT-C score in the discovery and replication samples. To adjust for significant global confounding factors, we followed a comprehensive analysis pipeline developed by Lehne *et al*. ^47^. The primary EWAS model used a DNAm µ>-value (the ratio of methylated probe intensity divided by the overall intensity) as the response variable and the continuous natural logarithm of PEth as the predictor variable of interest. Since previous studies have shown that a large number of CpGs were significantly associated with age ^50^, smoking status ^13^, race ^51^, HIV status and HIV-1 VL^12^, these variables were adjusted in the models. The cell proportions of 6 cell types were also adjusted in the models. The log10 of viral load (*log_10_ VL*) and ART adherence were adjusted in the replication sample. The same models were also used for EWAS on AUDIT-C score in discovery and replication samples, where the AUDIT-C score was a response variable. Epigenome-wide significance was set at a false discovery rate (FDR) < 0.05 in the discovery sample. Significance in the replication sample was set at a nominal *p* < 0.05.

1. *First generalized linear model* We performed a linear model to adjust for the confounders mentioned above in both the discovery and replication models. For discovery,

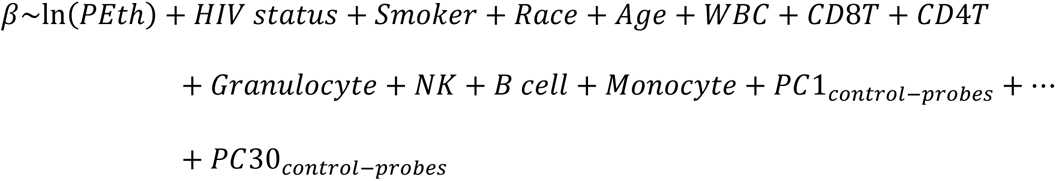

For replication,

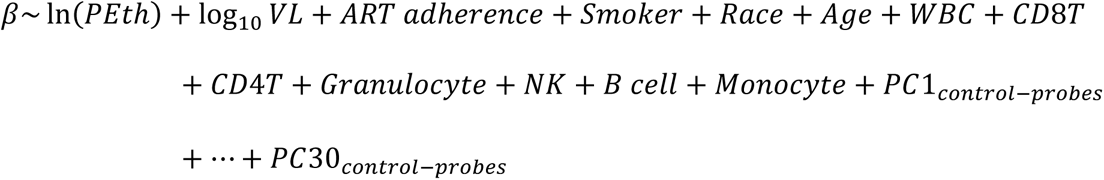
2. *Principal component analysis (PCA) of intermediary residuals* We then performed a PCA on the resulting regression residuals. The top five principal components on the residuals (PCs) (*PC1_residuals_,…, PC5_residuals_*) were adjusted in the final model.
3. *A final generalized linear model for identifying differential methylation* We performed a final generalized linear regression analysis for each methylation marker predicting the *β* as a function of the natural logarithm of the PEth value adjusted for technical and biological factors and the top 5 PC residuals derived from the model above.

### Meta-analysis of EWAS in Cohort 1

An EWAS meta-analysis was conducted by combining the discovery sample and the replication samples. For each CpG, we obtained effect size estimates and p-values from the two samples and weighted the effect size estimates by their estimated standard errors. Then, the summary statistics of the two samples were combined using a sample-size weighted meta-analysis using the METAL program ^52^. Epigenome-wide significance was set at a FDR <0.05.

### PolyGenic Methylation Score (PGMS)

We constructed a PGMS for each individual as a weighted sum of the individual CpG *µ>* values using the effect size estimated from the EWAS as weights ^14^. In detail, the PEth-related CpGs identified in the meta-analysis were chosen to construct the PGMS. Then, the PGMS was applied to establish a prediction model for HAD in Cohort 2.

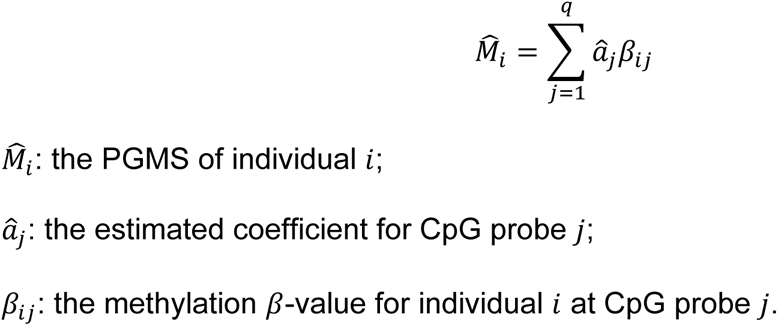

### Adjusted ***R^2^*** and incremental adjusted ***R^2^***

We used the adjusted *R^2^* to estimate the phenotypic variances explained by the DNA methylation. The adjusted *R^2^* accounted for the number of predictors in the model. The adjusted *R^2^*represented the percentage of variation explained by only the independent variables that affected the dependent variable. Here, the adjusted *R^2^* was the proportion of the variance of the PEth values or AUDIT-C scores that was explained by the PGMS or the individual CpG methylation.

We applied the incremental adjusted *R^2^* (incremental *R^2^*) as one of the parameters for feature selection as described below. The incremental *R^2^* was used to determine whether a new predictor increases the predictive ability above and beyond that provided by an existing model. It was calculated for each selected CpG or the linear combination of selected CpGs.

### Feature selection using elastic net regularization (ENR)

CpG features were separately preselected from the EWAS results on PEth and on AUDIT-C in Cohort 1. Using the ENR method, we performed a 10-fold cross-validation for feature selection in the training sample of Cohort 2. Here, we randomly selected 80% of the samples in Cohort 2 and cross-validated them to obtain the values for the ENR tuning parameters. The following steps were taken to select the CpG features and to evaluate their performance.

Step 1. *Preselection CpGs*. Because DNAm of CpGs under the epigenome-wide significance threshold may collectively account for phenotype variation and may improve prediction of a phenotype, we preselected PEth-associated CpGs with a meta *p* < 1E-04 from the meta-EWAS in Cohort 1 for both PEth and AUDIT-C. The preselected CpGs were used to establish the predictive model in the training set of Cohort 2.

Step 2. *Importance ranking CpGs*. In the training set of Cohort 2, we performed an ENR for feature selection among the preselected CpGs. We extracted the coefficients for the model with the lambda value corresponding to the minimum mean cross-validated error. This procedure was repeated *N* times. We excluded the CpGs with percentage of zero coefficients larger than 95%. All selected CpGs were reranked according to the summation of the absolute value of the *N* coefficients.

Step 3. *Model building by ENR in the training set*. CpG features were selected based on the area under the receiver operating characteristic curve (AUC), prediction accuracy, and the incremental *R^2^* for different numbers of CpG sets. The model with the best performance was determined, and the optimal values of the parameters in the best model were found by performing cross-validation in ENR.

Step 4. *Model performance testing in the testing set*. The performance of the CpG features selected from the training set were evaluated in the testing set using AUC, prediction accuracy, and the incremental *R^2^*.

All analyses were performed using R software (https://www.r-project.org/). ENR was performed using the function “cv.glmnet” in the “glmnet” package.

### Biological interpretation of the prediction model

Gene enrichment analysis was performed using the CpGs from the final prediction model to understand the underlying biological significance. We applied the web-accessible, gene annotation term-based Database for Annotation, Visualization and Integrated Discovery (DAVID) for gene enrichment analysis (http://david.niaid.nih.gov)^53^. The expanded DAVID Knowledgebase integrates almost all major and well-known public bioinformatics resources ^54^. A significant pathway was set as a nominal *p* < 1.00E-02.

## RESULTS

### EWAS identifies new DNA methylation CpGs for PEth but not for self-reported alcohol consumption

Two analyses of EWAS on PEth values and on the AUDIT-C scores were separately conducted in Cohort 1. Phenotypically, as expected, PEth level and AUDIT-C score were highly correlated (*r* = 0.45, *p* < 2.00E-16) (**Figure S1a**). Compared to the non-HAD group, the HAD group had a greater AUDIT-C score and a higher level of PEth (*p* = 3.47E-33) (**Figure S1b**).

In the discovery sample, the HAD group included more African Americans (AAs), had a higher rate of tobacco smokers and a lower level of ART adherence compared to the non-HAD group (*p* < 5.00E-02). In the replication sample, the prevalence of smoking in the HAD group was higher than in the non-HAD group, but this difference did not reach statistical significance (**Table 1**). Smoking status was still adjusted in the model to address potential smoking effects.

### Discovery EWAS on PEth and on AUDIT-C

We identified 9 epigenome-wide significant CpGs on PEth (FDR *p* = 1.22E-04∼4.68E-02) (**Figure S2a, Table S1**). The EWAS analysis showed minimal inflation (*λ* = 1.093) (**Figure S2b**). The 9 significant CpGs were located on 7 genes: *SLC7A11* (solute carrier family 7 member 11), *DYRK2* (dual specificity tyrosine phosphorylation regulated kinase 2), *FOXP1* (forkhead box P1), *SLC43A1* (solute carrier family 43 member 1), *WDR1* (WD repeat domain 1), *ABAT* (4-aminobutyrate aminotransferase), and *CCDC71* (coiled-coil domain containing 71). Six of 9 CpGs were negatively associated with PEth while 3 of 9 were positively associated with PEth.

We found no CpGs that reached an epigenome-wide significance threshold for self-reported AUDIT-C scores. Six of the 9 CpGs associated with PEth showed nominal association with AUDIT-C (nominal *p* ranged from 3.50E-03 to 4.76E-02): cg06690548 (*SLC7A11*), cg17962756, cg13442969 (*DYRK2*), cg11376147 (*SLC43A1*), cg00220102 (*ABAT*), and cg18590502 (*CCDC71*). It is noteworthy that all 9 CpGs associated with PEth showed the same direction as the associations with the AUDIT-C scores in the discovery set.

### Replication EWAS on PEth and on AUDIT-C scores

In the replication sample, we found 3 epigenome-wide significant CpGs associated with PEth: cg20414364 (*LOC728613*), cg10988872 (*LRCH4*), and cg01434144 (*STXBP5-AS1*) (FDR = 3.62E-02 ∼ 4.01E-02) (**Figure S3**). For the 9 PEth-associated CpGs identified in the discovery sample, we found that 6 of 9 CpGs showed nominal significance for PEth, although they did not reach epigenome-wide significance (nominal *p* ranged from 2.26E-06 to 2.80E-02) (**Table S1**). The 6 CpGs were located on 5 genes: cg06690548 (*SLC7A11*), cg17962756, cg13442969 (*DYRK2*), cg11376147 (*SLC43A1*), cg26689780 (*WDR1*), and cg18590502 (*CCDC71*).

As expected, the analysis of the EWAS on AUDIT-C scores revealed no CpG reaching epigenome-wide significance in the replication sample. Only 2 of 9 CpGs associated with PEth were nominally associated with AUDIT-C scores (cg11376147 in *SLC43A1* with *t* = -3.69 and *p* = 2.71E-04, cg26689780 in *WDR1* with *t* = 2.07 and *p* = 3.92E-02) and showed the same direction as the association of PEth.

### Meta-analysis

A meta-analysis revealed 102 epigenome-wide significant CpGs on PEth (FDR = 1.39E-06 ∼ 4.89E−02) (**Table 2** and **Figure 2a**). A majority of these CpGs (83 out of 102 CpGs) were in a gene region, including 24 CpGs in the promoter, 2 CpGs in the first exon, and 12 CpGs in the UTR regions. With a stringent significant threshold, 13 CpGs showed a Bonferroni adjusted *p* < 5.00E-02. These 13 CpGs mapped to 9 genes, including 6 novel genes for alcohol consumption (*DYRK2*, *PAK1*, *LOC728613*, *ATG7*, *TRA2B*, and *FBLN2*) and 3 genes (*SLC7A11*, *SLC43A1,* and *WDR1)* previously reported to be related to alcohol consumption ^29, 30, 55^.

**Figure 2.**
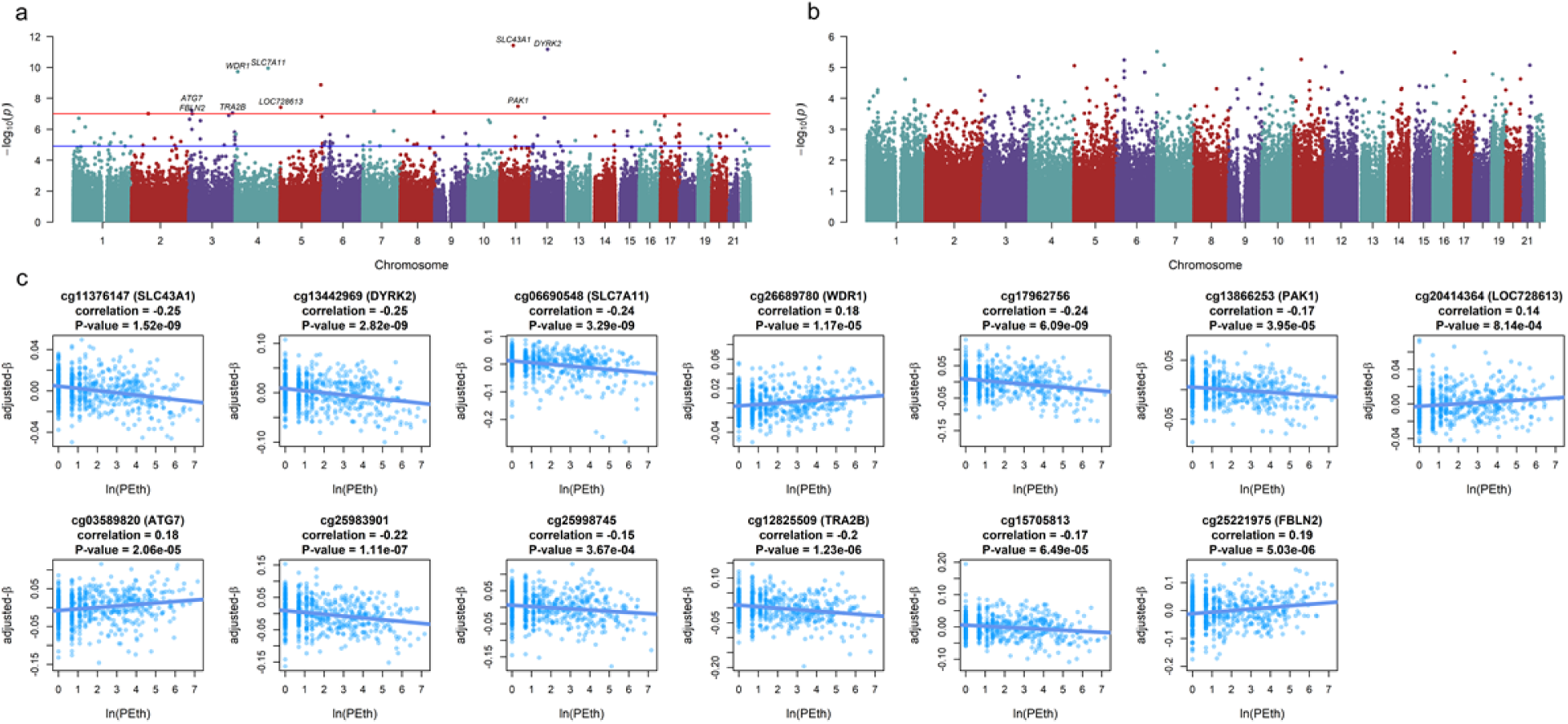
Meta-analysis of epigenome-wide association studies of alcohol consumption. **a.** Manhattan plot of chromosomal locations of − log_10_(*p*) for the association between Phosphatidylethanol (PEth) and 408,583 CpGs in the meta-analysis. **b.** Manhattan plot of chromosomal locations of − log_10_(*p*) for the association between Hazardous Alcohol Drinking (HAD) and 408,583 CpGs in the meta-analysis. The red line represents the threshold for Bonferroni-corrected p-value. The blue line represents the threshold for false discovery rate (FDR)-corrected p-value. **c.** Scatterplots of the adjusted *µ>*-values of the 13 CpGs against the natural logarithm of the PEth value. All 13 CpGs were significantly correlated with the natural logarithm of the PEth value with *p* < 1.00E-03.

**Table 2.** Significant epigenome-wide DNA methylation sites associated with Phosphatidylethanol (PEth) in meta-analysis of Cohort 1 (False Discovery Rate < 5.00E-02 in meta-analysis)

| Probe | | CHR | Position | Gene | Group | Incremental adjusted $R^2$ | Discovery | | Replication | | Meta-analysis | | | Reference |
| --- | --- | --- | --- | --- | --- | --- | --- | --- | --- | --- | --- | --- | --- | --- |
|  |  |  |  |  |  |  | t | P-value | t | P-value | Z-Score | P-value | FDR |  |
| 1 | cg11376147 | 11 | 57261198 | *SLC43A1* | Body | 4.31% | -5.14 | 3.90E-07 | -4.80 | 2.26E-06 | -6.95 | 3.79E-12 | 1.39E-06 | 29, 30 |
| 2 | cg13442969 | 12 | 68044208 | *DYRK2* | 5UTR | 4.94% | -5.52 | 5.34E-08 | -4.25 | 2.70E-05 | -6.86 | 6.81E-12 | 1.39E-06 | 30 |
| 3 | cg06690548 | 4 | 139162808 | *SLC7A11* | Body | 6.63% | -6.44 | 2.80E-10 | -2.59 | 1.01E-02 | -6.45 | 1.14E-10 | 1.55E-05 | 29, 30 |
| 4 | cg26689780 | 4 | 10079554 | *WDR1* | Body | 4.30% | 5.12 | 4.29E-07 | 3.90 | 1.13E-04 | 6.37 | 1.89E-10 | 1.94E-05 | 29 |
| 5 | cg17962756 | 5 | 172769199 | NA | NA | 5.42% | -5.83 | 9.51E-09 | -2.70 | 7.34E-03 | -6.06 | 1.35E-09 | 1.11E-04 | 29, 30 |
| 6 | cg13866253 | 11 | 77093001 | *PAK1* | Body | 2.16% | -3.70 | 2.43E-04 | -4.28 | 2.41E-05 | -5.53 | 3.30E-08 | 2.22E-03 |  |
| 7 | cg20414364 | 5 | 1608614 | *LOC728613* | Body | 0.94% | 2.49 | 1.30E-02 | 5.59 | 4.28E-08 | 5.50 | 3.80E-08 | 2.22E-03 |  |
| 8 | cg03589820 | 3 | 11585825 | *ATG7* | Body | 3.39% | 4.56 | 6.35E-06 | 3.08 | 2.23E-03 | 5.42 | 5.89E-08 | 2.96E-03 |  |
| 9 | cg25983901 | 7 | 46972700 | NA | NA | 2.98% | -4.34 | 1.68E-05 | -3.36 | 8.69E-04 | -5.40 | 6.80E-08 | 2.96E-03 | 29 |
| 10 | cg25998745 | 8 | 142028625 | NA | NA | 2.45% | -3.89 | 1.11E-04 | -3.79 | 1.78E-04 | -5.38 | 7.24E-08 | 2.96E-03 | 30, 60 |
| 11 | cg12825509 | 3 | 185648568 | *TRA2B* | Body | 3.34% | -4.55 | 6.68E-06 | -3.01 | 2.76E-03 | -5.35 | 8.70E-08 | 3.09E-03 | 30, 38 |
| 12 | cg15705813 | 2 | 70297499 | NA | NA | 2.54% | -3.94 | 9.21E-05 | -3.65 | 3.03E-04 | -5.33 | 9.83E-08 | 3.09E-03 | 29, 30 |
| 13 | cg25221975 | 3 | 13663444 | *FBLN2* | Body | 3.72% | 4.82 | 1.88E-06 | 2.69 | 7.41E-03 | 5.33 | 9.83E-08 | 3.09E-03 | 30 |
| 14 | cg19825437 | 3 | 169383292 | *NA* | NA | 3.71% | -4.76 | 2.47E-06 | -2.63 | 8.85E-03 | -5.28 | 1.27E-07 | 3.70E-03 | 29, 30 |
| 15 | cg27376514 | 17 | 17058422 | *MPRIP* | Body | 2.82% | 4.17 | 3.53E-05 | 3.29 | 1.11E-03 | 5.26 | 1.40E-07 | 3.81E-03 |  |
| 16 | cg19731612 | 5 | 176559334 | *NSD1* | TSS1500 | 1.88% | -3.45 | 6.02E-04 | -4.10 | 4.95E-05 | -5.25 | 1.53E-07 | 3.91E-03 | 29, 30 |
| 17 | cg02583484 | 12 | 54677008 | *HNRNPA1;*<br>*HNRPA1L-2* | Body | 2.25% | -3.74 | 2.06E-04 | -3.71 | 2.41E-04 | -5.22 | 1.82E-07 | 4.37E-03 | 29, 30, 60 |
| 18 | cg07167185 | 1 | 24120017 | *LYPLA2* | Body | 2.44% | 3.23 | 1.33E-03 | 4.24 | 2.82E-05 | 5.20 | 1.99E-07 | 4.52E-03 |  |
| 19 | cg00294109 | 3 | 3219781 | *CRBN* | Body | 2% | 3.53 | 4.61E-04 | 3.87 | 1.26E-04 | 5.17 | 2.28E-07 | 4.91E-03 |  |
| 20 | cg24351003 | 10 | 88013210 | *GRID1* | Body | 2.33% | -3.82 | 1.50E-04 | -3.53 | 4.61E-04 | -5.16 | 2.53E-07 | 5.17E-03 |  |
| 21 | cg18590502 | 3 | 49203081 | *CCDC71* | 5UTR | 3.99% | -4.96 | 9.64E-07 | -2.21 | 2.80E-02 | -5.14 | 2.79E-07 | 5.42E-03 | 30 |
| 22 | cg00944421 | 16 | 68269483 | *ESRP2* | Body | 2.97% | -4.26 | 2.41E-05 | -2.92 | 3.72E-03 | -5.11 | 3.28E-07 | 6.09E-03 |  |
| 23 | cg24238409 | 10 | 93998677 | *CPEB3* | Body | 3.37% | 4.50 | 8.31E-06 | 2.58 | 1.03E-02 | 5.09 | 3.64E-07 | 6.46E-03 |  |
| 24 | cg02256576 | 16 | 66995192 | *CES3* | 5UTR | 2.50% | -3.93 | 9.78E-05 | -3.20 | 1.46E-03 | -5.04 | 4.71E-07 | 7.96E-03 | 29, 30 |
| 25 | cg19869698 | 17 | 80058686 | *NA* | NA | 3.64% | 4.74 | 2.72E-06 | 2.29 | 2.24E-02 | 5.03 | 4.87E-07 | 7.96E-03 |  |
| 26 | cg08250921 | 16 | 88111009 | *NA* | NA | 3.67% | 4.73 | 2.83E-06 | 2.24 | 2.54E-02 | 5.01 | 5.33E-07 | 8.38E-03 |  |
| 27 | cg23090529 | 1 | 51442133 | *NA* | NA | 2.41% | -3.87 | 1.24E-04 | -3.16 | 1.71E-03 | -4.96 | 7.10E-07 | 1.07E-02 | 30, 60 |
| 28 | cg23482898 | 3 | 12858887 | CAND2 | Body | 2.01% | 3.58 | 3.75E-04 | 3.42 | 6.96E-04 | 4.89 | 1.00E-06 | 1.46E-02 | 30 |
| 29 | cg01425762 | 16 | 81666633 | CMIP | Body | 1.03% | 2.61 | 9.30E-03 | 4.48 | 9.91E-06 | 4.87 | 1.11E-06 | 1.56E-02 |  |
| 30 | cg11704631 | 21 | 36395663 | RUNX1 | Body | 2.74% | -4.13 | 4.25E-05 | -2.73 | 6.59E-03 | -4.86 | 1.15E-06 | 1.57E-02 | 30 |
| 31 | cg08616943 | 7 | 130552600 | NA | NA | 2.11% | -3.64 | 3.00E-04 | -3.26 | 1.23E-03 | -4.85 | 1.26E-06 | 1.66E-02 |  |
| 32 | cg23028286 | 15 | 51614521 | CYP19A1 | 5UTR | 3.39% | -4.58 | 5.72E-06 | -2.17 | 3.04E-02 | -4.84 | 1.32E-06 | 1.66E-02 |  |
| 33 | cg21550372 | 14 | 100908908 | WDR25 | Body | 2.56% | -3.97 | 8.12E-05 | -2.83 | 4.87E-03 | -4.83 | 1.35E-06 | 1.66E-02 |  |
| 34 | cg13966547 | 1 | 2406284 | PLCH2 | TSS1500 | 1.36% | -2.96 | 3.19E-03 | -4.00 | 7.47E-05 | -4.83 | 1.38E-06 | 1.66E-02 |  |
| 35 | cg00220102 | 16 | 8806756 | ABAT | TSS200 | 4.12% | -5.02 | 6.98E-07 | -1.62 | 1.06E-01 | -4.81 | 1.49E-06 | 1.74E-02 |  |
| 36 | cg00166216 | 3 | 194407860 | FAM43A | 1stExon | 2.40% | -3.87 | 1.23E-04 | -2.93 | 3.60E-03 | -4.81 | 1.54E-06 | 1.75E-02 |  |
| 37 | cg03044573 | 1 | 173835265 | GAS5;<br>SNORD78;<br>SNORD44;<br>SNORD80;<br>SNORD79 | TSS1500 | 1.53% | -3.14 | 1.80E-03 | -3.73 | 2.22E-04 | -4.77 | 1.81E-06 | 1.96E-02 | 30 |
| 38 | cg14395885 | 9 | 130700923 | DPM2 | TSS200 | 0.78% | -2.32 | 2.05E-02 | -4.66 | 4.43E-06 | -4.77 | 1.82E-06 | 1.96E-02 |  |
| 39 | cg24135793 | 19 | 13122567 | NFIX | Body | 1.92% | -3.48 | 5.53E-04 | -3.31 | 1.01E-03 | -4.77 | 1.88E-06 | 1.96E-02 | 30, 38 |
| 40 | cg06925984 | 17 | 77767242 | NA | NA | 2.02% | -3.56 | 4.07E-04 | -3.21 | 1.45E-03 | -4.76 | 1.92E-06 | 1.96E-02 |  |
| 41 | cg03840289 | 4 | 2262318 | MXD4 | Body | 1.85% | 3.42 | 6.83E-04 | 3.36 | 8.54E-04 | 4.75 | 1.99E-06 | 1.98E-02 |  |
| 42 | cg11846968 | 20 | 31823545 | *PLUNC* | TSS1500 | 0.90% | -2.50 | 1.28E-02 | -4.45 | 1.12E-05 | -4.75 | 2.03E-06 | 1.98E-02 |  |
| 43 | cg10692140 | 6 | 30496072 | *NA* | NA | 1.54% | -3.18 | 1.54E-03 | -3.66 | 2.91E-04 | -4.74 | 2.15E-06 | 2.05E-02 |  |
| 44 | cg23747342 | 12 | 25539794 | *NA* | NA | 1.06% | 2.65 | 8.40E-03 | 4.17 | 3.73E-05 | 4.71 | 2.54E-06 | 2.27E-02 |  |
| 45 | cg05303280 | 15 | 51632611 | *GLDN* | TSS1500 | 3.77% | -4.85 | 1.65E-06 | -1.69 | 9.23E-02 | -4.70 | 2.58E-06 | 2.27E-02 |  |
| 46 | cg10891521 | 17 | 81047941 | *METRNL* | Body | 0.94% | 2.53 | 1.18E-02 | 4.32 | 2.00E-05 | 4.70 | 2.63E-06 | 2.27E-02 |  |
| 47 | cg27477373 | 19 | 56879645 | *ZNF542* | TSS200 | 1.88% | -3.45 | 6.08E-04 | -3.25 | 1.27E-03 | -4.70 | 2.66E-06 | 2.27E-02 |  |
| 48 | cg17521665 | 6 | 106546704 | *PRDM1* | TSS200 | 1.56% | -3.15 | 1.71E-03 | -3.55 | 4.25E-04 | -4.69 | 2.76E-06 | 2.27E-02 |  |
| 49 | cg13548452 | 14 | 22573606 | *NA* | NA | 2.06% | -3.62 | 3.18E-04 | -3.05 | 2.42E-03 | -4.69 | 2.77E-06 | 2.27E-02 |  |
| 50 | cg23352030 | 20 | 62198469 | *PRIC285* | 1stExon | 1.31% | 2.95 | 3.36E-03 | 3.84 | 1.43E-04 | 4.69 | 2.78E-06 | 2.27E-02 |  |
| 51 | cg13610455 | 20 | 37054900 | *LOC388796;*<br>*SNORA71B* | TSS1500 | 1.43% | 3.03 | 2.56E-03 | 3.69 | 2.52E-04 | 4.68 | 2.84E-06 | 2.27E-02 |  |
| 52 | cg06059663 | 1 | 245319431 | *KIF26B* | Body | 2.19% | -3.67 | 2.72E-04 | -2.92 | 3.74E-03 | -4.67 | 2.93E-06 | 2.29E-02 |  |
| 53 | cg21845080 | 3 | 196065306 | *TM4SF19* | TSS200 | 0.51% | 1.95 | 5.14E-02 | 4.93 | 1.22E-06 | 4.67 | 3.01E-06 | 2.29E-02 |  |
| 54 | cg09801924 | 11 | 65425948 | *RELA* | Body | 1.40% | 3.01 | 2.73E-03 | 3.70 | 2.42E-04 | 4.67 | 3.08E-06 | 2.29E-02 |  |
| 55 | cg00970435 | 17 | 66380327 | *ARSG* | Body | 1.17% | -2.78 | 5.62E-03 | -3.98 | 8.36E-05 | -4.67 | 3.08E-06 | 2.29E-02 |  |
| 56 | cg23579062 | 9 | 34457500 | *C9orf25;*<br>*DNAI1* | TSS1500 | 1.50% | -3.06 | 2.34E-03 | -3.58 | 3.88E-04 | -4.66 | 3.20E-06 | 2.34E-02 | 29, 30 |
| 57 | cg04202267 | 2 | 169431900 | LASS6 | Body | 2.38% | 3.81 | 1.56E-04 | 2.71 | 7.01E-03 | 4.65 | 3.33E-06 | 2.39E-02 |  |
| 58 | cg20970380 | 1 | 116676103 | C1orf161 | 3UTR | 3.76% | -4.80 | 2.04E-06 | -1.59 | 1.12E-01 | -4.63 | 3.61E-06 | 2.54E-02 |  |
| 59 | cg00717678 | 17 | 1554577 | PRPF8;<br>RILP | TSS1500 | 3.33% | 4.54 | 7.13E-06 | 1.90 | 5.79E-02 | 4.63 | 3.70E-06 | 2.56E-02 |  |
| 60 | cg14817906 | 2 | 97466833 | CNNM4 | Body | 2.73% | 4.08 | 5.14E-05 | 2.36 | 1.89E-02 | 4.62 | 3.92E-06 | 2.63E-02 |  |
| 61 | cg27653384 | 22 | 22293118 | PPM1F | Body | 0.71% | 2.26 | 2.42E-02 | 4.51 | 8.62E-06 | 4.62 | 3.93E-06 | 2.63E-02 | 30 |
| 62 | cg22537604 | 19 | 43857074 | CD177 | TSS1500 | 3.14% | -4.38 | 1.43E-05 | -2.02 | 4.37E-02 | -4.61 | 4.01E-06 | 2.64E-02 |  |
| 63 | cg20732160 | 3 | 48590040 | PFKFB4 | Body | 2.62% | -4.02 | 6.68E-05 | -2.42 | 1.59E-02 | -4.60 | 4.24E-06 | 2.75E-02 |  |
| 64 | cg01883662 | 3 | 196065289 | TM4SF19 | TSS200 | 0.57% | 2.04 | 4.14E-02 | 4.67 | 4.16E-06 | 4.57 | 4.80E-06 | 3.07E-02 |  |
| 65 | cg24366564 | 17 | 2843149 | RAP1GAP2 | Body | 1.58% | 3.17 | 1.63E-03 | 3.34 | 9.26E-04 | 4.56 | 5.13E-06 | 3.18E-02 |  |
| 66 | cg03394159 | 8 | 29197844 | DUSP4 | Body | 2.02% | 3.60 | 3.50E-04 | 2.89 | 4.02E-03 | 4.56 | 5.13E-06 | 3.18E-02 |  |
| 67 | cg06906869 | 13 | 52734154 | NEK3 | TSS1500 | 1.10% | 2.72 | 6.81E-03 | 3.87 | 1.27E-04 | 4.55 | 5.38E-06 | 3.28E-02 |  |
| 68 | cg10440877 | 2 | 208378475 | NA | NA | 1.54% | -3.14 | 1.78E-03 | -3.32 | 9.84E-04 | -4.52 | 6.13E-06 | 3.64E-02 |  |
| 69 | cg17840178 | 6 | 30709803 | FLOT1 | Body | 2.78% | -4.14 | 4.05E-05 | -2.17 | 3.05E-02 | -4.52 | 6.15E-06 | 3.64E-02 |  |
| 70 | cg00966482 | 6 | 11111926 | HERV-FRD;<br>LOC221710 | 5UTR | 3.87% | 4.90 | 1.26E-06 | 1.33 | 1.84E-01 | 4.51 | 6.43E-06 | 3.74E-02 |  |
| 71 | cg15033653 | 12 | 113587581 | CCDC42B | TSS200 | 2.77% | 4.11 | 4.55E-05 | 2.16 | 3.17E-02 | 4.51 | 6.50E-06 | 3.74E-02 |  |
| 72 | cg09635954 | 7 | 29605624 | *PRR15* | 5UTR | 0.79% | -2.35 | 1.93E-02 | -4.23 | 2.98E-05 | -4.50 | 6.72E-06 | 3.82E-02 |  |
| 73 | cg26841068 | 1 | 203456691 | *PRELP* | 3UTR | 2.76% | -4.16 | 3.70E-05 | -2.15 | 3.24E-02 | -4.50 | 6.83E-06 | 3.82E-02 | 29, 38 |
| 74 | cg19939130 | 1 | 158978468 | *IFI16* | TSS1500 | 2.24% | -3.78 | 1.73E-04 | -2.58 | 1.01E-02 | -4.49 | 7.03E-06 | 3.88E-02 |  |
| 75 | cg14728380 | 17 | 80280330 | *SECTM1* | Body | 2.11% | 3.66 | 2.79E-04 | 2.68 | 7.58E-03 | 4.48 | 7.51E-06 | 4.01E-02 |  |
| 76 | cg06983052 | 1 | 90288099 | *LRRC8D* | 5UTR | 2.02% | -3.56 | 4.03E-04 | -2.77 | 5.90E-03 | -4.48 | 7.54E-06 | 4.01E-02 | 29, 30 |
| 77 | cg24136754 | 22 | 37403978 | *C22orf33* | TSS200 | 1.76% | 3.31 | 9.93E-04 | 3.03 | 2.64E-03 | 4.48 | 7.59E-06 | 4.01E-02 |  |
| 78 | cg21366673 | 6 | 30459512 | *HLA-E* | Body | 3.37% | -4.59 | 5.68E-06 | -1.63 | 1.03E-01 | -4.47 | 7.66E-06 | 4.01E-02 | 29 |
| 79 | cg23598378 | 6 | 42072986 | *C6orf132* | Body | 0.95% | 2.53 | 1.17E-02 | 3.95 | 9.34E-05 | 4.47 | 7.83E-06 | 4.05E-02 | 29 |
| 80 | cg11826008 | 20 | 34249284 | *CPNE1;*<br>*RBM12* | 5UTR | 2.13% | -3.68 | 2.56E-04 | -2.65 | 8.43E-03 | -4.46 | 8.03E-06 | 4.10E-02 | 38 |
| 81 | cg24136700 | 17 | 17696044 | *RAI1* | 5UTR | 0.59% | 2.07 | 3.92E-02 | 4.46 | 1.10E-05 | 4.46 | 8.30E-06 | 4.15E-02 |  |
| 82 | cg06937549 | 5 | 179046350 | *HNRNPH1* | Body | 2.07% | -3.61 | 3.33E-04 | -2.70 | 7.34E-03 | -4.46 | 8.32E-06 | 4.15E-02 |  |
| 83 | cg20699548 | 8 | 71060638 | *NCOA2* | Body | 2.48% | -3.96 | 8.64E-05 | -2.30 | 2.21E-02 | -4.44 | 8.83E-06 | 4.35E-02 |  |
| 84 | cg18568145 | 1 | 155225764 | *FAM189B* | TSS1500 | 2.51% | -3.96 | 8.55E-05 | -2.27 | 2.39E-02 | -4.44 | 9.01E-06 | 4.38E-02 |  |
| 85 | cg11302401 | 6 | 6688847 | *NA* | NA | 2.45% | -3.96 | 8.42E-05 | -2.32 | 2.10E-02 | -4.43 | 9.26E-06 | 4.45E-02 |  |
| 86 | cg13706315 | 9 | 134724316 | *NA* | NA | 0.92% | 2.51 | 1.25E-02 | 3.91 | 1.08E-04 | 4.43 | 9.57E-06 | 4.47E-02 | 30 |
| 87 | cg22503354 | 12 | 7341644 | *PEX5* | TSS1500 | 2.69% | 4.06 | 5.64E-05 | 2.10 | 3.64E-02 | 4.43 | 9.59E-06 | 4.47E-02 |  |
| 88 | cg22496559 | 3 | 196065318 | *TM4SF19* | TSS200 | 1.08% | 2.65 | 8.26E-03 | 3.71 | 2.42E-04 | 4.42 | 9.71E-06 | 4.47E-02 |  |
| 89 | cg23684449 | 16 | 46919194 | *GPT2* | 5UTR | 1.48% | -3.08 | 2.17E-03 | -3.23 | 1.33E-03 | -4.42 | 9.73E-06 | 4.47E-02 | 30, 38 |
| 90 | cg23144445 | 8 | 57471162 | NA | NA | 2.52% | -3.97 | 8.29E-05 | -2.23 | 2.64E-02 | -4.42 | 9.88E-06 | 4.48E-02 |  |
| 91 | cg17206604 | 3 | 150088779 | NA | NA | 1.52% | 3.09 | 2.10E-03 | 3.17 | 1.67E-03 | 4.41 | 1.03E-05 | 4.60E-02 |  |
| 92 | cg10447615 | 1 | 109506805 | *CLCC1* | TSS1500 | 2.34% | 3.79 | 1.69E-04 | 2.37 | 1.83E-02 | 4.41 | 1.04E-05 | 4.60E-02 |  |
| 93 | cg19536127 | 2 | 47404286 | *CALM2* | TSS1500 | 1.86% | 3.40 | 7.14E-04 | 2.81 | 5.28E-03 | 4.40 | 1.07E-05 | 4.62E-02 |  |
| 94 | cg01304182 | 16 | 30409908 | *ZNF48* | Body | 1.19% | 2.80 | 5.29E-03 | 3.53 | 4.66E-04 | 4.40 | 1.08E-05 | 4.62E-02 |  |
| 95 | cg02003183 | 14 | 103415882 | *CDC42BPB* | Body | 2.03% | 3.55 | 4.21E-04 | 2.64 | 8.65E-03 | 4.40 | 1.09E-05 | 4.62E-02 | 30, 38, 60 |
| 96 | cg22994830 | 7 | 623846 | *PRKAR1B* | Body | 2.10% | 3.62 | 3.23E-04 | 2.57 | 1.06E-02 | 4.40 | 1.09E-05 | 4.62E-02 | 29 |
| 97 | cg22274745 | 2 | 182451537 | *CERKL* | Body | 1.42% | 3.01 | 2.75E-03 | 3.26 | 1.23E-03 | 4.40 | 1.10E-05 | 4.62E-02 |  |
| 98 | cg27155460 | 10 | 45420821 | *TMEM72* | Body | 2.71% | 4.11 | 4.70E-05 | 2.03 | 4.29E-02 | 4.39 | 1.11E-05 | 4.65E-02 |  |
| 99 | cg14718379 | 7 | 71806067 | *CALN1* | Body | 0.97% | -2.55 | 1.10E-02 | -3.78 | 1.80E-04 | -4.38 | 1.17E-05 | 4.85E-02 |  |
| 100 | cg03329019 | 1 | 221051117 | NA | NA | 1.93% | -3.48 | 5.42E-04 | -2.70 | 7.27E-03 | -4.38 | 1.20E-05 | 4.87E-02 |  |
| 101 | cg11599718 | 12 | 123357128 | *VPS37B* | Body | 1.24% | 2.85 | 4.52E-03 | 3.44 | 6.46E-04 | 4.38 | 1.20E-05 | 4.87E-02 |  |
| 102 | cg01211396 | 6 | 30624478 | *DHX16* | Body | 1.54% | 3.13 | 1.82E-03 | 3.10 | 2.10E-03 | 4.37 | 1.22E-05 | 4.89E-02 | 29 |

Interestingly, even with an increased sample size in the meta-analysis, we found no epigenome-wide significant CpG site of the meta-EWAS on AUDIT-C scores (**Figure 2b**).

We further tested the correlation between the *µ>*-values of the 13 CpGs with Bonferroni significance and PEth. All 13 CpGs were significantly correlated with PEth levels after the model was adjusted for confounding factors (**Figure 2c**), four of the 13 CpGs were positively correlated with PEth, and the remaining 9 CpGs were negatively correlated with PEth.

### PEth-associated CpG sites improves the prediction of HAD in Cohort 1

Because PEth itself was highly correlated with AUDIT-C scores and differed significantly between the HAD and the non-HAD groups, we were interested in whether PEth-associated CpG DNAm improved the prediction of HAD compared to the prediction of HAD using PEth alone. We found that the AUC was 74.2% for PEth alone, 76.9% with the 13 Bonferroni significant CpGs and PEth, and 88.3% with the 102 epigenome-wide significant CpGs and PEth (**Figure S4**). Thus, DNAm features improved the prediction of hazardous alcohol consumption compared to PEth alone in the same cohort.

### PGMS derived from 102 PEth-associated CpGs is correlated with alcohol consumption in an independent sample

To be consistent with the analysis in Cohort 1, we performed an EWAS on AUDIT-C score in Cohort 2. We found no epigenome-wide significant CpG for AUDIT-C. An EWAS for a full scale of AUDIT score also revealed no significant CpG.

We found that a PGMS constructed from the 102 PEth-associated CpGs was highly correlated with the self-reported 10-item AUDIT score in Cohort 2 (*r* = 0.40, *p* < 8.09E-20). The incremental *R^2^* of the association between the PGMS corresponding to 102 PEth-related CpGs and the full AUDIT score was 0.1002, which implied that the PGMS explained 10.02% of the variance of the full AUDIT score in an independent population (**Figure S5a**).

We further tested whether the PGMS derived from the PEth-associated CpGs was separately correlated with self-reported alcohol consumption (AUDIT-C, first 3-items of AUDIT) and self-reported problem alcohol drinking behaviors (AUDIT-P, item 4-10 of full AUDIT). We found that the PGMS was significantly correlated with AUDIT-C score (*r* = 0.37, *p* = 4.60E-16) (**Figure S5b**) and AUDIT-P score (*r* = 0.35, *p* = 3.70E-11) (**Figure S5c**). The correlation of the PGMS was slightly stronger with the AUDIT-C score than with the AUDIT-P score.

### PEth-associated DNA methylation CpG sites predict HAD in Cohort 2

Notably, we found no statistically significant difference in the characteristics between the training set and the testing set in Cohort 2 (**Table S2**). Using the ENR method, we preselected PEth-associated CpGs with meta *p* < 1E-04 from the meta-EWAS in Cohort 1. A total of 302 CpGs were preselected to build a predictive model in the training set of Cohort 2. After excluding the CpGs with a percentage of zero coefficients larger than 95% using ENR, a total of 249 CpGs remained for model construction. All 249 CpGs were ranked according to the summation of the absolute value of the *N* coefficients. As shown in **Figure 3a**, a panel of 130 CpGs showed the greatest AUC with 91.31% and the highest incremental *R^2^* with 21.29% in the training set. Therefore, a model derived from these 130 CpGs was validated in the testing set.

**Figure 3.**
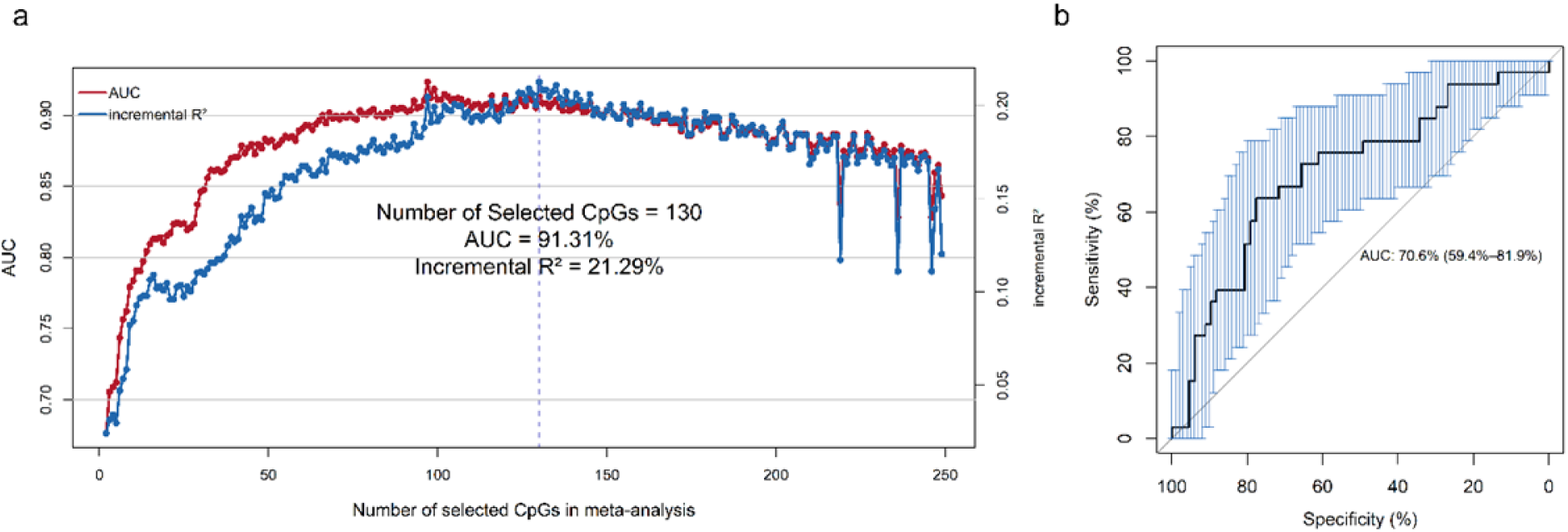
Feature selection using elastic net regularization (ENR). **a.** The area under the receiver operating characteristic curve (AUC) and the incremental adjusted *R^2^* (incremental *R^2^*) of the selected CpG sites using ENR method (pre-selection CpG sites cutoff *p* < 1.00E-04 in the training set of Cohort 2). Incremental *R^2^* denotes the difference in adjusted *R^2^* between the model with the predicted variable and the model without the predicted variable. **b.** The receiver operating characteristic (ROC) curve for Hazardous Alcohol Drinking (HAD) prediction in the testing set of Cohort 2 using 130 out of 302 CpGs with *p* < 1E-04 in the meta-analysis.

In the testing set, we found that the model with the 130 CpGs showed an AUC of 70.60%, a balanced accuracy of 60.00%, and an incremental *R^2^* of 3.79% (**Figure 3b**). The results show that the 130 selected PEth-associated CpGs enabled the good prediction of HAD. Notably, the panel of 130 CpGs included 48 epigenome-wide significant CpGs for meta-EWAS on PEth in Cohort 1. We summarized the information of the 130 selected CpGs in **Table S3**.

Using the same approach for the analysis of feature selection of AUDIT-C-associated CpGs from Cohort 1 to predict HAD in Cohort 2, a panel of 18 CpGs were selected from 54 CpGs with *p* < 1E-04. In the training set, the AUC was 70.13%, and the incremental *R^2^* was 2.18%. In the testing set, the AUC was 57.6% (46.1%-69.1%), and the incremental *R^2^* was 1.07%.

### Biological interpretation of the 130 identified PEth-associated CpGs

The 130 CpGs from the final predictive model were annotated on 111 genes. Gene enrichment analysis using the 111 genes yielded 28 significant annotation terms (*p* < 1.00E-02, **Figure S6**). The top significant pathways included GO:0048519∼negative regulation of biological process (p=2.63E-05); GO:0048523∼negative regulation of cellular process (*p* = 1.80E-04; GO:0030155∼regulation of cell adhesion (*p* = 1.00E-03); GO:0010605∼negative regulation of macromolecule metabolic process (*p* = 2.00E-03); GO:0009892∼negative regulation of metabolic process (*p* = 2.00E-03); GO:0010629∼negative regulation of gene expression (*p* = 3.00E-03); GO:0065009∼regulation of molecular function (*p* = 3.00E-03), and GO:0044093∼positive regulation of molecular function (*p* = 3.00E-03).

## DISCUSSION

Using samples from two distinct populations, we have demonstrated that an objective phenotype, PEth, is a robust phenotype for identifying DNAm in blood associated with HAD and that PEth-associated CpGs are predictive of HAD. We revealed 102 CpGs associated with PEth, while none of the CpGs were associated with self-reported alcohol consumption. A PGMS derived from the 102 CpGs explained 10.02% of the variance of alcohol consumption in a demographically and clinically independent sample. We further showed that the 102 CpGs combined with PEth improved 14% of AUC of predicting HAD compared to the AUC of predicting HAD by PEth alone. Importantly, we identified a panel of 130 CpGs, that were relevant to PEth levels in a mostly HIV-positive sample and that predicted self-reported HAD in an HIV-negative sample. The 130 CpGs included 18 CpGs that were previously included in the DNAm biomarker panel for prediction of alcohol consumption by Liu *et al*. ^30^ However, a panel of CpGs related to self-reported AUDIT-C score showed poor predictive performance for HAD. Together, these findings suggest that PEth-associated DNAm features, but not DNAm for self-reported alcohol consumption, is a robust biomarker to predict hazardous alcohol consumption that may have potential clinical utility.

Emerging evidence suggests that a set of epigenetic modification markers across different tissues is more stable and reproducible than we previously expected ^56^. In this study, we replicated 32 CpGs that had previously reported associations with alcohol consumption or alcohol use disorders. For example, three promoter CpGs, cg19731612 on *NSD1* (FDR = 3.91E-03) ^29, 30^, cg03044573 on *SNORD78* (FDR = 1.96E-02) ^30^, and cg23579062 on *DNAI1* (FDR = 2.34E-02) ^29, 30^ that were associated with alcohol consumption in previous studies were also significantly associated with PEth in our study. In addition, we revealed multiple new PEth-associated CpGs that are located on the genes involved in tyrosine autophosphorylation, catalyzed phosphorylation of histones H3 and H2B (*DYRK2*) and the serine/threonine p21-activating kinases (*PAK1*), modulation of the p53-dependent cell cycle pathways during prolonged metabolic stress (*ATG7*), sequence-specific serine/arginine splicing factor (*TRA2B*) functions, and extracellular matrix protein (*FBLN2*). These results suggest that alcohol consumption alters DNA methylation on the genes involved in the cellular process and epigenetic programming. Although the findings do not elucidate the etiology of alcohol drinking behavior in brain, the significant CpGs suggest a peripheral mechanism of how alcohol consumption changes the epigenome in peripheral cells, which may lead to alcohol use-related medical disorders.

The 102 PEth-associated CpGs identified in a mostly HIV-positive population collectively explained 10.02% of the variance of HAD in an HIV-negative population, suggesting the stability of the DNAm effect of alcohol exposure. Notably, the 10.02% effect size of the PGMS in our study is comparable with the previously reported 12%∼13.8% effect size of a PGMS in a study with a 10-fold larger sample size (N = 13,317) than this study^30^. We further showed that PGMS was not only significantly associated with recent alcohol consumption (AUDIT-C score) (*r* = 0.37, *p* = 4.60E-16) but was strongly associated with the problematic consequences of alcohol use (AUDIT-P score) (*r* = 0.35, *p* = 3.70E-11), further indicating that DNAm is a relatively stable marker for the long-term effects of alcohol consumption. Future studies evaluating DNAm marker stability for alcohol consumption using longitudinal DNAm measurements are needed.

The reproducible CpGs suggest a robust, consistent epigenetic response to alcohol consumption that can serve as biomarkers for clinical use. Using a machine learning approach, we identified a set of 130 CpGs that enables the distinction of HAD and non-HAD individuals. One of the common challenges for machine learning prediction is model overfitting. We took several steps to address this concern: 1) feature preselection and selection were conducted in two different cohorts; 2) the processes of feature selection and model evaluation were carried out in the same cohort but in different sets without overlapping samples; and 3) we applied a newly developed machine learning ENR method to select features in a combination of 10-fold cross-validation. Compared to two traditional penalized regression methods, Ridge ^57^ and the least absolute shrinkage and selection operator (LASSO) ^58^, ENR has the advantage of selecting informative features without compromising predictive accuracy and has been shown to outperform both the Ridge and LASSO methods ^59^. With these strengths of the analytical approach, we showed that a panel of 130 CpGs performed fairly well with an AUC of 70.60%, a balanced accuracy of 60.00%, and an incremental *R^2^* of 3.79% in the testing sample set. Although the AUC in our study was less than the previously reported AUC of 0.90-0.99 with 144 CpGs ^30^, our result is less likely to be inflated because of our analytical approach to avoid data overfitting. In the previous study, the model building and evaluation were performed using the same sample set while we performed the prediction analysis in the training and testing set separately.

Several limitations should be considered in interpreting the current findings. 1) There was a lack of power to detect sex-specific associations between CpGs and HAD. It is well known that HAD in men and women is epidemiologically and mechanistically different. The individuals in Cohort 1 were all men and approximately 50% of the individuals in Cohort 2 were women. These samples are insufficient to seek sex-specific DNAm markers. 2) The DNAm signatures were identified from whole blood samples that lacked cell-type specific profiles. Future analyses using cell-type-specific CpGs may improve the prediction performance. 3) The 130 CpGs in the DNAm signature were preselected from an HIV-positive sample, while the prediction model was built and validated in an HIV-negative sample. We expect to improve the predictive efficiency in a relatively homogenous sample in future studies. 4) Validation of the prediction panel on other alcohol use-related phenotypes, e.g., alcohol use disorder, is necessary to confidently claim the predictive performance and accuracy for clinical use.

In summary, to the best of our knowledge, this is the first study to demonstrate that PEth is a robust phenotype for detecting subtle DNAm changes associated with alcohol consumption compared to self-reported alcohol use data. PEth-associated DNAm markers predicted HAD with a good accuracy. These findings suggest that DNAm signatures may have clinical utility as biomarkers for alcohol consumption, and further development and testing of these biomarkers are warranted.

## ACKNOWLEDGMENTS

The authors appreciate the support of the Veterans Aging Study Cohort Biomarker Core and the Yale Center of Genomic Analysis.

## AVAILABILITY OF DATA AND MATERIALS

Demographic variables, clinical variables and methylation status for the VACS samples were submitted to the GEO dataset (GSE117861) and are available to the public. All codes for analysis are also available upon a request to the corresponding author.

## AUTHORS’ CONTRIBUTIONS

XL was responsible for the bioinformatics data processing and statistical analysis. ACJ provided DNA samples and clinical data and contributed to the interpretation of findings and manuscript preparation. KS contributed to the manuscript preparation. JHK contributed to the interpretation of findings and manuscript preparation. RS provided DNA samples and clinical data and contributed to manuscript preparation. KX was responsible for the study design, study protocol, sample preparation, data analysis, interpretation of findings, and manuscript preparation. All authors read and approved the final manuscript.

## Supplemental Tables and Figures

**Table S1.** Significant epigenome-wide DNA methylation sites associated with Phosphatidylethanol (PEth) in discovery of Cohort 1 (False Discovery Rate < 5E-02 in discovery of Cohort 1)

**Table S2.** Demographic and clinical characteristics for the feature selection set (training) and validation set (testing) in Cohort 2

**Table S3.** The 130 selected CpGs for predicting Hazardous Alcohol Drinking (HAD) using elastic net regularization (ENR)

**Figure S1.** Correlation between Phosphatidylethanol (PEth) and Alcohol Use Disorders Identification Test-Consumption items (AUDIT-C) score in Cohort 1. **a.** Scatter plot showing significant association between the ln(PEth) value and the AUDIT-C score (The Pearson correlation between ln(PEth) and AUDIT-C is 0.45 (95% CI: 0.39, 0.51) with *p* < 2.00E-16). **b.** Violin plot showing significant difference of the ln(PEth) value between non-Hazardous Alcohol Drinking (non-HAD) (ADUIT-C >= 4) participants and HAD participants. The P-value of two sample t-test for non-HAD and HAD is 3.47E-33, which indicates that the biomarker PEth and alcohol consumption are significantly correlated.

**Figure S2.** Manhattan plot and quantile-quantile (QQ) plot for the discovery set of Cohort 1. **a.** Manhattan plot of the chromosomal locations of − log_10_(*p*) for the epigenome-wide association in 437,722 CpGs among the 580 males in the discovery sample set. The red line represents the threshold for Bonferroni-corrected p-value. The blue line represents the threshold for false discovery rate (FDR)-corrected p-value. **b.** QQ plot for association at all 437,722 CpGs. *λ = 1.093* in the discovery epigenome-wide association analysis.

**Figure S3.** Manhattan plot and quantile-quantile (QQ) plot for the replication set of Cohort 1. **a.** Manhattan plot of the chromosomal locations of − log_10_(*p*) for the epigenome-wide association in 846,604 CpGs among the 467 males in the replication sample set. The red line represents the threshold for Bonferroni-corrected p-value. The blue line represents the threshold for false discovery rate (FDR)-corrected p-value. **b.** QQ plot for the association at all 846,604 CpGs. *λ = 1.146* in the replication epigenome-wide association analysis.

**Figure S4.** Receiver Operating Characteristic (ROC) curve for predicting Hazardous Alcohol Drinking (HAD). ROC curve for predicting HAD by Phosphatidylethanol (PEth) alone, PEth with 13 CpGs (Bonferroni corrected p-value less than 5.00E-02), and PEth with 102 CpGs (false discovery rate (FDR)-corrected p-value less than 5.00E-02) for samples in Cohort 1.

**Figure S5.** Scatterplots for alcohol-related phenotype vs. PolyGenic Methylation Score (PGMS) constructed by 102 Phosphatidylethanol (PEth)-related CpGs. **a.** Scatterplots of Alcohol Use Disorders Identification Test (AUDIT) score vs. PGMS. **b.** Scatterplots of Alcohol Use Disorders Identification Test-Consumption items (AUDIT-C) vs. PGMS. **c.** Scatterplots of Alcohol Use Disorders Identification Test-Problem items (AUDIT-P) score vs. PGMS.

**Figure S6.** Database for Annotation, Visualization and Integrated Discovery (DAVID) pathway analysis for the 130 CpGs selected by elastic net regularization (ENR).

