## Supplementary Information for "DNA Methylation Signature on Phosphatidylethanol, not Self-Reported Alcohol Consumption, Predicts Hazardous Alcohol Consumption in Two Distinct Populations"

[1]Department of Psychiatry, Yale School of Medicine, New Haven, CT, USA; [2]VA Connecticut Healthcare System, West Haven, CT, USA; [3]Yale University School of Medicine, New Haven Veterans Affairs Connecticut Healthcare System, New Haven, CT, USA; [4]Boston University School of Medicine, Boston, MA, USA; [5]Child Study Center, Yale School of Medicine, New Haven, CT, USA; [6]Department of Neuroscience, Yale School of Medicine, New Haven, CT, USA

**All corresponding to**

Ke Xu, MD, PhD

Associate Professor of Psychiatry

Yale School of Medicine

**Table S1.** Significant epigenome-wide DNA methylation sites associated with Phosphatidylethanol (PEth) in discovery of Cohort 1 (False Discovery Rate < 5E-02 in discovery of Cohort 1)

| Probe | | CHR | Position | Gene | Group | Incremental adjusted $R^2$ | Discovery | | | Replication | |
|---|---|---|---|---|---|---|---|---|---|---|---|
| | | | | | | | t | p | FDR | t | p |
| 1 | cg06690548 | 4 | 139162808 | *SLC7A11* | Body | 6.62% | -6.44 | 2.80E-10 | 1.22E-04 | -2.59 | 1.01E-02 |
| 2 | cg17962756 | 5 | 172769199 | *NA* | NA | 5.49% | -5.83 | 9.51E-09 | 2.08E-03 | -2.70 | 7.34E-03 |
| 3 | cg13442969 | 12 | 68044208 | *DYRK2* | 5UTR | 4.93% | -5.52 | 5.34E-08 | 7.77E-03 | -4.25 | 2.70E-05 |
| 4 | cg20525486 | 3 | 71111923 | *FOXP1* | Body | 4.37% | 5.20 | 2.92E-07 | 2.78E-02 | NA | NA |
| 5 | cg11376147 | 11 | 57261198 | *SLC43A1* | Body | 4.28% | -5.14 | 3.90E-07 | 2.78E-02 | -4.80 | 2.26E-06 |
| 6 | cg26689780 | 4 | 10079554 | *WDR1* | Body | 4.25% | 5.12 | 4.29E-07 | 2.78E-02 | 3.90 | 1.13E-04 |
| 7 | cg04304130 | 6 | 11111894 | *LOC221710; HERV-FRD* | 5UTR | 4.24% | 5.11 | 4.45E-07 | 2.78E-02 | NA | NA |
| 8 | cg00220102 | 16 | 8806756 | *ABAT* | TSS200 | 4.09% | -5.02 | 6.98E-07 | 3.81E-02 | -1.62 | 1.06E-01 |
| 9 | cg18590502 | 3 | 49203081 | *CCDC71* | 5UTR | 3.99% | -4.96 | 9.64E-07 | 4.68E-02 | -2.21 | 2.80E-02 |

**Table S2.** Demographic and clinical characteristics for the feature selection set (training) and validation set (testing) in Cohort 2

| | Training Set (N = 402) | Testing Set (N = 100) |
|---|---|---|
| AUDIT | 5.42 ± 5.83 | 6.81 ± 7.12 |
| Age (year) | 28.93 ± 9.01 | 28.72 ± 7.92 |
| Sex (male, %) | 43.28 | 47.00 |
| Race (AA, %) | 18.91 | 21.00 |
| Smoker (%) | 21.64 | 17.00 |
| CD4+ T (%) | 0.18 ± 0.05 | 0.18 ± 0.05 |
| CD8+ T (%) | 0.09 ± 0.04 | 0.09 ± 0.04 |
| NK (%)[a] | 0.03 ± 0.03 | 0.04 ± 0.04 |
| B cell (%)[a] | 0.07 ± 0.03 | 0.07 ± 0.02 |
| Monocyte (%)[a] | 0.08 ± 0.02 | 0.08 ± 0.02 |
| Granulocyte (%)[a] | 0.59 ± 0.09 | 0.59 ± 0.08 |

AA: African American, AUDIT: Alcohol Use Disorders Identification Test

[a]Cell type compositions estimated by methylation

**Table S3.** The 130 selected CpGs for predicting Hazardous Alcohol Drinking (HAD) using Elastic Net Regularization

| Probe | | CHR | Position | Gene | Group | Incremental adjusted $R^2$ | Z-Score | P-value | FDR | Reference |
|---|---|---|---|---|---|---|---|---|---|---|
| 1 | cg06690548 | 4 | 139162808 | *SLC7A11* | Body | 6.63% | -6.45 | 1.14E-10 | 1.55E-05 | Liu et al., 2018; Wilson et al., 2019 |
| 2 | cg15049370 | 5 | 149186389 | *PPARGC1B* | Body | 0.72% | 4.09 | 4.29E-05 | 9.24E-02 | |
| 3 | cg00716016 | 9 | 3180374 | *NA* | NA | 1.45% | -4.07 | 4.81E-05 | 9.77E-02 | |
| 4 | cg09191335 | 20 | 35241157 | *SLA2* | 3UTR | 2.09% | 4.33 | 1.51E-05 | 5.51E-02 | |
| 5 | cg19825437 | 3 | 169383292 | *NA* | NA | 3.71% | -5.28 | 1.27E-07 | 3.70E-03 | Liu et al., 2018; Wilson et al., 2019 |
| 6 | cg25221975 | 3 | 13663444 | *FBLN2* | Body | 3.72% | 5.33 | 9.83E-08 | 3.09E-03 | Liu et al., 2018 |
| 7 | cg08688548 | 15 | 83317398 | *CPEB1* | TSS1500 | 2.31% | -4.08 | 4.58E-05 | 9.44E-02 | |
| 8 | cg13121938 | 13 | 114871497 | *RASA3* | Body | 1.68% | 4.11 | 3.97E-05 | 8.97E-02 | |
| 9 | cg26856289 | 1 | 24307516 | *SFRS13A* | TSS1500 | 1.92% | -3.92 | 8.76E-05 | 1.29E-01 | |
| 10 | cg13706315 | 9 | 134724316 | *NA* | NA | 0.92% | 4.43 | 9.57E-06 | 4.47E-02 | Liu et al., 2018 |
| 11 | cg23728471 | 17 | 42345689 | *SLC4A1* | TSS200 | 1.64% | 4.03 | 5.64E-05 | 1.04E-01 | |
| 12 | cg26103104 | 5 | 139490895 | *NA* | NA | 1.20% | -4.21 | 2.53E-05 | 7.61E-02 | Liu et al., 2018 |
| 13 | cg17415265 | 17 | 63225111 | *NA* | NA | 0.58% | 4.04 | 5.25E-05 | 1.01E-01 | |

| | | | | | | | | | | |
|---|---|---|---|---|---|---|---|---|---|---|
| 14 | cg09635954 | 7 | 29605624 | *PRR15* | 5UTR | 0.79% | -4.5 | 6.72E-06 | 3.82E-02 | |
| 15 | cg00717678 | 17 | 1554577 | *PRPF8; RILP* | TSS1500 | 3.33% | 4.63 | 3.70E-06 | 2.56E-02 | |
| 16 | cg18076842 | 11 | 93262607 | *C11orf75* | 5UTR | 1.89% | -4.3 | 1.70E-05 | 5.78E-02 | |
| 17 | cg02256576 | 16 | 66995192 | *CES3* | 5UTR | 2.50% | -5.04 | 4.71E-07 | 7.96E-03 | Liu et al., 2018; Wilson et al., 2019 |
| 18 | cg24713122 | 11 | 46389164 | *DGKZ* | Body | 0.77% | 4.27 | 1.99E-05 | 6.52E-02 | |
| 19 | cg22007110 | 7 | 30737599 | *NA* | NA | 0.58% | 4.01 | 6.13E-05 | 1.09E-01 | Wilson et al., 2019 |
| 20 | cg06230839 | 21 | 43189892 | *NA* | NA | 1.71% | -3.91 | 9.30E-05 | 1.30E-01 | |
| 21 | cg09801924 | 11 | 65425948 | *RELA* | Body | 1.40% | 4.67 | 3.08E-06 | 2.29E-02 | |
| 22 | cg20732160 | 3 | 48590040 | *PFKFB4* | Body | 2.62% | -4.6 | 4.24E-06 | 2.75E-02 | |
| 23 | cg13442969 | 12 | 68044208 | *DYRK2* | 5UTR | 4.94% | -6.86 | 6.81E-12 | 1.39E-06 | Liu et al., 2018 |
| 24 | cg24135793 | 19 | 13122567 | *NFIX* | Body | 1.92% | -4.77 | 1.88E-06 | 1.96E-02 | Liu et al., 2018 |
| 25 | cg01278873 | 19 | 44764101 | *ZNF233* | 5UTR | 1.23% | -4.35 | 1.39E-05 | 5.21E-02 | |
| 26 | cg00256932 | 22 | 51041732 | *MAPK8IP2* | 1stExon | 2.32% | 4.28 | 1.91E-05 | 6.29E-02 | |
| 27 | cg05095466 | 5 | 37207622 | *C5orf42* | Body | 1.87% | -3.97 | 7.32E-05 | 1.18E-01 | |
| 28 | cg19098763 | 1 | 50513661 | *ELAVL4* | TSS200 | 1.55% | -3.92 | 9.05E-05 | 1.30E-01 | |

| | | | | | | | | | | |
|---|---|---|---|---|---|---|---|---|---|---|
| 29 | cg23028286 | 15 | 51614521 | *CYP19A1* | 5UTR | 3.39% | -4.84 | 1.32E-06 | 1.66E-02 | |
| 30 | cg21090033 | 3 | 196065357 | *TM4SF19* | TSS200 | 0.81% | 4.03 | 5.54E-05 | 1.04E-01 | |
| 31 | cg16663980 | 1 | 158809049 | *MNDA* | 5UTR | 3.56% | -4.01 | 6.11E-05 | 1.09E-01 | |
| 32 | cg01005506 | 10 | 64565768 | *ADO* | 1stExon | 1.42% | -4.13 | 3.60E-05 | 8.66E-02 | |
| 33 | cg10381071 | 15 | 70391035 | *TLE3* | TSS1500 | 0.56% | -4.34 | 1.41E-05 | 5.23E-02 | |
| 34 | cg06937549 | 5 | 179046350 | *HNRNPH1* | Body | 2.07% | -4.46 | 8.32E-06 | 4.15E-02 | |
| 35 | cg02538135 | 19 | 2200125 | *DOT1L* | Body | 0.67% | 4.14 | 3.53E-05 | 8.66E-02 | |
| 36 | cg01881182 | 16 | 8806531 | *ABAT* | TSS1500 | 2.48% | -4.1 | 4.14E-05 | 9.04E-02 | |
| 37 | cg01883662 | 3 | 196065289 | *TM4SF19* | TSS200 | 0.57% | 4.57 | 4.80E-06 | 3.07E-02 | |
| 38 | cg13341668 | 3 | 50359909 | *HYAL2* | TSS1500 | 1.53% | -4 | 6.22E-05 | 1.09E-01 | |
| 39 | cg03546163 | 6 | 35654363 | *FKBP5* | 5UTR | 1.00% | -4.19 | 2.84E-05 | 7.72E-02 | |
| 40 | cg19623438 | 1 | 149230829 | *NA* | NA | 1.89% | -4.25 | 2.17E-05 | 6.87E-02 | |
| 41 | cg19731612 | 5 | 176559334 | *NSD1* | TSS1500 | 1.88% | -5.25 | 1.53E-07 | 3.91E-03 | Liu et al., 2018; Wilson et al., 2019 |
| 42 | cg21550372 | 14 | 100908908 | *WDR25* | Body | 2.56% | -4.83 | 1.35E-06 | 1.66E-02 | |
| 43 | cg07387591 | 20 | 17208649 | *PCSK2* | Body | 0.82% | 4.03 | 5.56E-05 | 1.04E-01 | |

| | | | | | | | | | | |
|---|---|---|---|---|---|---|---|---|---|---|
| 44 | cg08908135 | 13 | 113405377 | *ATP11A* | Body | 1.51% | 3.94 | 8.08E-05 | 1.25E-01 | |
| 45 | cg05303280 | 15 | 51632611 | *GLDN* | TSS1500 | 3.77% | -4.7 | 2.58E-06 | 2.27E-02 | |
| 46 | cg24136700 | 17 | 17696044 | *RAI1* | 5UTR | 0.59% | 4.46 | 8.30E-06 | 4.15E-02 | |
| 47 | cg27653384 | 22 | 22293118 | *PPM1F* | Body | 0.71% | 4.62 | 3.93E-06 | 2.63E-02 | Liu et al., 2018 |
| 48 | cg26468430 | 7 | 1572571 | *MAFK* | 5UTR | 0.53% | 4.05 | 5.01E-05 | 9.92E-02 | |
| 49 | cg02605760 | 7 | 72723152 | *NSUN5* | TSS1500 | 0.95% | 3.91 | 9.13E-05 | 1.30E-01 | |
| 50 | cg22566142 | 5 | 96298493 | *LNPEP* | 5UTR | 2.79% | -3.94 | 8.03E-05 | 1.25E-01 | Wilson et al., 2019 |
| 51 | cg03163545 | 16 | 1593415 | *IFT140; TMEM204* | Body | 1.80% | 4.28 | 1.89E-05 | 6.26E-02 | |
| 52 | cg03458172 | 3 | 157815672 | *SHOX2* | 3UTR | 0.98% | -4.03 | 5.69E-05 | 1.05E-01 | |
| 53 | cg27477373 | 19 | 56879645 | *ZNF542* | TSS200 | 1.88% | -4.7 | 2.66E-06 | 2.27E-02 | |
| 54 | cg07866212 | 8 | 23836952 | *NA* | NA | 1.57% | 4.09 | 4.32E-05 | 9.24E-02 | |
| 55 | cg03523740 | 1 | 32645027 | *TXLNA* | TSS1500 | 1.78% | -4.35 | 1.34E-05 | 5.21E-02 | Liu et al., 2018; Wilson et al., 2019 |
| 56 | cg14014731 | 9 | 19378679 | *RPS6* | Body | 1.08% | -3.92 | 8.79E-05 | 1.29E-01 | |
| 57 | cg23352030 | 20 | 62198469 | *PRIC285* | 1stExon | 1.31% | 4.69 | 2.78E-06 | 2.27E-02 | |
| 58 | cg01156249 | 16 | 4714794 | *MGRN1* | Body | 3.80% | 4.23 | 2.36E-05 | 7.30E-02 | |

| | | | | | | | | | |
|---|---|---|---|---|---|---|---|---|---|
| 59 | cg22537604 | 19 | 43857074 | *CD177* | TSS1500 | 3.14% | -4.61 | 4.01E-06 | 2.64E-02 |
| 60 | cg20283107 | 8 | 124788969 | *FAM91A1* | Body | 2.79% | -4.31 | 1.66E-05 | 5.74E-02 |
| 61 | cg10891521 | 17 | 81047941 | *METRNL* | Body | 0.94% | 4.7 | 2.63E-06 | 2.27E-02 |
| 62 | cg14718379 | 7 | 71806067 | *CALN1* | Body | 0.97% | -4.38 | 1.17E-05 | 4.85E-02 |
| 63 | cg25746394 | 19 | 45450501 | *APOC2* | 5UTR | 1.52% | 4.2 | 2.70E-05 | 7.68E-02 |
| 64 | cg17953300 | 11 | 65418265 | *SIPA1* | 3UTR | 1.54% | 4.35 | 1.34E-05 | 5.21E-02 |
| 65 | cg10603800 | 17 | 80408909 | *C17orf62* | TSS1500 | 1.13% | 3.97 | 7.15E-05 | 1.16E-01 |
| 66 | cg07769421 | 17 | 80816851 | *TBCD* | Body | 1.60% | 4.33 | 1.53E-05 | 5.52E-02 |
| 67 | cg25673668 | 8 | 895601 | *NA* | NA | 0.92% | -4.09 | 4.35E-05 | 9.25E-02 |
| 68 | cg08404225 | 3 | 3151899 | *IL5RA* | 5UTR | 1.19% | 3.93 | 8.45E-05 | 1.27E-01 |
| 69 | cg27155460 | 10 | 45420821 | *TMEM72* | Body | 2.71% | 4.39 | 1.11E-05 | 4.65E-02 |
| 70 | cg03401875 | 5 | 178684658 | *ADAMTS2* | Body | 0.48% | -4.13 | 3.67E-05 | 8.66E-02 |
| 71 | cg23291200 | 19 | 1473179 | *APC2* | 3UTR | 2.17% | 3.94 | 8.03E-05 | 1.25E-01 |
| 72 | cg11302401 | 6 | 6688847 | *NA* | NA | 2.45% | -4.43 | 9.26E-06 | 4.45E-02 |
| 73 | cg11494699 | 11 | 36588217 | *RAG1* | TSS1500 | 1.22% | 3.94 | 8.31E-05 | 1.26E-01 |

| | | | | | | | | | | |
|---|---|---|---|---|---|---|---|---|---|---|
| 74 | cg01281718 | 6 | 71376634 | *SMAP1* | TSS1500 | 0.76% | 4.08 | 4.57E-05 | 9.44E-02 | Liu et al., 2018 |
| 75 | cg01182455 | 9 | 131313189 | *NA* | NA | 2.71% | 4.1 | 4.06E-05 | 9.00E-02 | |
| 76 | cg07714319 | 2 | 16822974 | *FAM49A* | 5UTR | 0.57% | -4 | 6.26E-05 | 1.09E-01 | |
| 77 | cg20785674 | 1 | 150123490 | *PLEKHO1* | Body | 2.28% | 3.91 | 9.28E-05 | 1.30E-01 | |
| 78 | cg20699548 | 8 | 71060638 | *NCOA2* | Body | 2.48% | -4.44 | 8.83E-06 | 4.35E-02 | |
| 79 | cg15326297 | 2 | 235464680 | *NA* | NA | 2.28% | 4.19 | 2.76E-05 | 7.72E-02 | |
| 80 | cg15033653 | 12 | 113587581 | *CCDC42B* | TSS200 | 2.77% | 4.51 | 6.50E-06 | 3.74E-02 | |
| 81 | cg15636519 | 2 | 191894418 | *STAT4* | 3UTR | 1.48% | -3.98 | 6.92E-05 | 1.14E-01 | |
| 82 | cg26841068 | 1 | 203456691 | *PRELP* | 3UTR | 2.76% | -4.5 | 6.83E-06 | 3.82E-02 | Wilson et al., 2019 |
| 83 | cg18121224 | 5 | 176559563 | *NSD1* | TSS1500 | 2.11% | -4.35 | 1.38E-05 | 5.21E-02 | Wilson et al., 2019 |
| 84 | cg06644515 | 1 | 173834831 | *SNORD47; GAS5; SNORD81; SNORD80; SNORD78; SNORD79* | TSS1500 | 1.39% | -3.92 | 8.80E-05 | 1.29E-01 | Liu et al., 2018; Wilson et al., 2019 |
| 85 | cg27458485 | 12 | 99139571 | *ANKS1B* | Body | 2.28% | 4.1 | 4.07E-05 | 9.00E-02 | |
| 86 | cg21194066 | 11 | 67052509 | *ADRBK1* | Body | 2.21% | -4.35 | 1.34E-05 | 5.21E-02 | |

| | | | | | | | | | | |
|---|---|---|---|---|---|---|---|---|---|---|
| 87 | cg09482421 | 6 | 27470541 | *NA* | NA | 0.71% | 4.11 | 4.00E-05 | 8.98E-02 | |
| 88 | cg12825509 | 3 | 185648568 | *TRA2B* | Body | 3.34% | -5.35 | 8.70E-08 | 3.09E-03 | Liu et al., 2018 |
| 89 | cg03598938 | 2 | 191502738 | *NA* | NA | 1.60% | -4.06 | 4.99E-05 | 9.92E-02 | |
| 90 | cg13966547 | 1 | 2406284 | *PLCH2* | TSS1500 | 1.36% | -4.83 | 1.38E-06 | 1.66E-02 | |
| 91 | cg11599718 | 12 | 123357128 | *VPS37B* | Body | 1.24% | 4.38 | 1.20E-05 | 4.87E-02 | |
| 92 | cg08119655 | 11 | 7595925 | *PPFIBP2* | Body | 1.85% | -4.16 | 3.14E-05 | 8.12E-02 | |
| 93 | cg00294109 | 3 | 3219781 | *CRBN* | Body | 2.00% | 5.17 | 2.28E-07 | 4.91E-03 | |
| 94 | cg10035272 | 6 | 142467043 | *VTA1* | TSS1500 | 1.77% | -4.07 | 4.80E-05 | 9.77E-02 | |
| 95 | cg17243654 | 1 | 109816280 | *CELSR2* | Body | 2.49% | 4.08 | 4.50E-05 | 9.37E-02 | |
| 96 | cg23090529 | 1 | 51442133 | *NA* | NA | 2.41% | -4.96 | 7.10E-07 | 1.07E-02 | Liu et al., 2018; Philibert et al., 2018 |
| 97 | cg15995125 | 1 | 17746597 | *RCC2* | Body | 0.88% | 4.31 | 1.62E-05 | 5.74E-02 | |
| 98 | cg26037936 | 11 | 120417671 | *NA* | NA | 0.85% | -4.05 | 5.09E-05 | 9.96E-02 | |
| 99 | cg03404339 | 12 | 52639319 | *KRT7* | Body | 1.12% | -4.18 | 2.94E-05 | 7.82E-02 | |
| 100 | cg15081033 | 3 | 140805567 | *SPSB4* | Body | 1.80% | 4.02 | 5.73E-05 | 1.05E-01 | |
| 101 | cg14259466 | 10 | 135090997 | *ADAM8* | TSS1500 | 0.95% | 3.98 | 6.86E-05 | 1.14E-01 | |

| | | | | | | | | | | |
|---|---|---|---|---|---|---|---|---|---|---|
| 102 | cg14395885 | 9 | 130700923 | *DPM2* | TSS200 | 0.78% | -4.77 | 1.82E-06 | 1.96E-02 | |
| 103 | cg23747342 | 12 | 25539794 | *NA* | NA | 1.06% | 4.71 | 2.54E-06 | 2.27E-02 | |
| 104 | cg16423756 | 11 | 122526190 | *UBASH3B* | TSS1500 | 3.02% | 4.31 | 1.64E-05 | 5.74E-02 | |
| 105 | cg04506190 | 3 | 129323941 | *PLXND1* | Body | 1.69% | -3.92 | 8.85E-05 | 1.29E-01 | |
| 106 | cg14107488 | 4 | 7033722 | *TBC1D14* | 3UTR | 0.80% | 4 | 6.21E-05 | 1.09E-01 | |
| 107 | cg10803871 | 5 | 174082675 | *NA* | NA | 1.62% | -4.18 | 2.95E-05 | 7.82E-02 | |
| 108 | cg01628467 | 17 | 7037406 | *NA* | NA | 1.36% | -4.19 | 2.78E-05 | 7.72E-02 | |
| 109 | cg08250921 | 16 | 88111009 | *NA* | NA | 3.67% | 5.01 | 5.33E-07 | 8.38E-03 | |
| 110 | cg10447615 | 1 | 109506805 | *CLCC1* | TSS1500 | 2.34% | 4.41 | 1.04E-05 | 4.60E-02 | |
| 111 | cg22215453 | 19 | 55852665 | *SUV420H2* | 5UTR | 0.77% | 4.12 | 3.70E-05 | 8.66E-02 | |
| 112 | cg11704631 | 21 | 36395663 | *RUNX1* | Body | 2.74% | -4.86 | 1.15E-06 | 1.57E-02 | Liu et al., 2018 |
| 113 | cg21786227 | 1 | 154951605 | *CKS1B* | Body | 2.51% | -3.93 | 8.49E-05 | 1.27E-01 | |
| 114 | cg22994830 | 7 | 623846 | *PRKAR1B* | Body | 2.10% | 4.4 | 1.09E-05 | 4.62E-02 | Wilson et al., 2019 |
| 115 | cg19939130 | 1 | 158978468 | *IFI16* | TSS1500 | 2.24% | -4.49 | 7.03E-06 | 3.88E-02 | |
| 116 | cg03840289 | 4 | 2262318 | *MXD4* | Body | 1.85% | 4.75 | 1.99E-06 | 1.98E-02 | |

| | | | | | | | | | | |
|---|---|---|---|---|---|---|---|---|---|---|
| 117 | cg18068637 | 5 | 168245286 | *SLIT3* | Body | 1.04% | -4.16 | 3.20E-05 | 8.16E-02 | |
| 118 | cg08204159 | 19 | 54642290 | *CNOT3* | 5UTR | 1.28% | -4.05 | 5.15E-05 | 9.98E-02 | |
| 119 | cg19056833 | 13 | 114749079 | *RASA3* | Body | 2.70% | 4.04 | 5.46E-05 | 1.03E-01 | |
| 120 | cg08893087 | 1 | 93416982 | *FAM69A* | Body | 1.92% | -3.98 | 6.91E-05 | 1.14E-01 | |
| 121 | cg15613761 | 19 | 46404710 | *MYPOP* | Body | 2.30% | -4.21 | 2.57E-05 | 7.62E-02 | |
| 122 | cg15705813 | 2 | 70297499 | *NA* | NA | 2.54% | -5.33 | 9.83E-08 | 3.09E-03 | Liu et al., 2018; Wilson et al., 2019 |
| 123 | cg06059663 | 1 | 245319431 | *KIF26B* | Body | 2.19% | -4.67 | 2.93E-06 | 2.29E-02 | |
| 124 | cg11899596 | 8 | 145663791 | *NFKBIL2* | Body | 0.52% | 4.15 | 3.39E-05 | 8.49E-02 | |
| 125 | cg10361922 | 17 | 40925790 | *VPS25* | Body | 1.54% | -3.99 | 6.68E-05 | 1.14E-01 | |
| 126 | cg17206604 | 3 | 150088779 | *NA* | NA | 1.52% | 4.41 | 1.03E-05 | 4.60E-02 | |
| 127 | cg22503354 | 12 | 7341644 | *PEX5* | TSS1500 | 2.69% | 4.43 | 9.59E-06 | 4.47E-02 | |
| 128 | cg19536127 | 2 | 47404286 | *CALM2* | TSS1500 | 1.86% | 4.4 | 1.07E-05 | 4.62E-02 | |
| 129 | cg00407659 | 5 | 150538414 | *ANXA6* | TSS1500 | 1.06% | -4.03 | 5.62E-05 | 1.04E-01 | Wilson et al., 2019 |
| 130 | cg18590502 | 3 | 49203081 | *CCDC71* | 5UTR | 3.99% | -5.14 | 2.79E-07 | 5.42E-03 | Liu et al., 2018 |

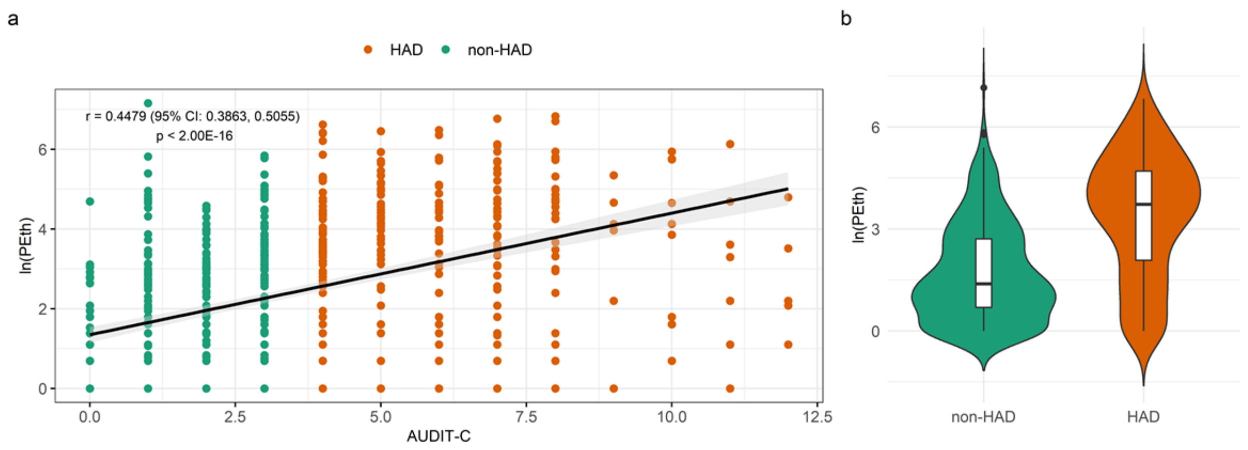

**Figure S1.** Correlation between Phosphatidylethanol (PEth) and Alcohol Use Disorders Identification Test-Consumption items (AUDIT-C) score in Cohort 1. **a.** Scatter plot showing significant association between the ln(PEth) value and the AUDIT-C score (The Pearson correlation between ln(PEth) and AUDIT-C is 0.45 (95% CI: 0.39, 0.51) with $p$ < 2.00E-16). **b.** Violin plot showing significant difference of the ln(PEth) value between non-Hazardous Alcohol Drinking (non-HAD) (ADUIT-C >= 4) participants and HAD participants. The P-value of two sample t-test for non-HAD and HAD is 3.47E-33, which indicates that the biomarker PEth and alcohol consumption are significantly correlated.

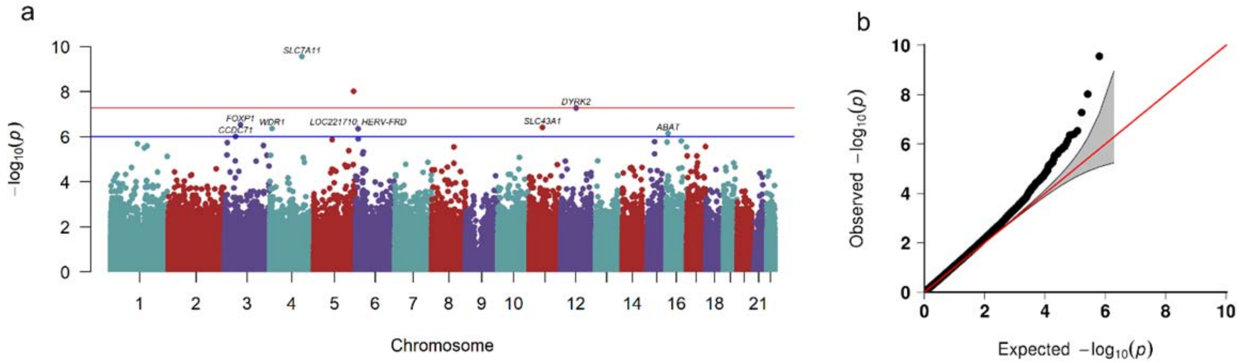

**Figure S2**. Manhattan plot and quantile-quantile (QQ) plot for the discovery set of Cohort 1. **a.** Manhattan plot of the chromosomal locations of $-\log_{10}(p)$ for the epigenome-wide association in 437,722 CpGs among the 580 males in the discovery sample set. The red line represents the threshold for Bonferroni-corrected p-value. The blue line represents the threshold for false discovery rate (FDR)-corrected p-value. **b.** QQ plot for association at all 437,722 CpGs. $\lambda = 1.093$ in the discovery epigenome-wide association analysis.

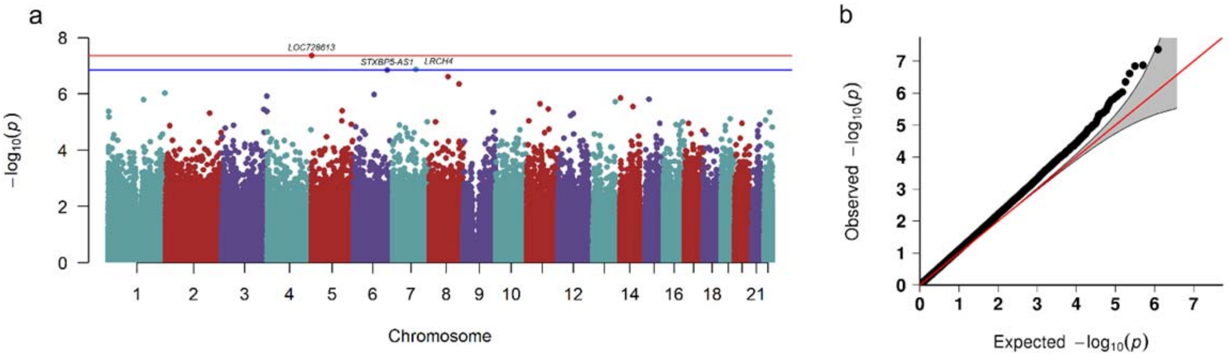

**Figure S3**. Manhattan plot and quantile-quantile (QQ) plot for the replication set of Cohort 1. **a.** Manhattan plot of the chromosomal locations of $-\log_{10}(p)$ for the epigenome-wide association in 846,604 CpGs among the 467 males in the replication sample set. The red line represents the threshold for Bonferroni-corrected p-value. The blue line represents the threshold for false discovery rate (FDR)-corrected p-value. **b.** QQ plot for the association at all 846,604 CpGs. $\lambda = 1.146$ in the replication epigenome-wide association analysis.

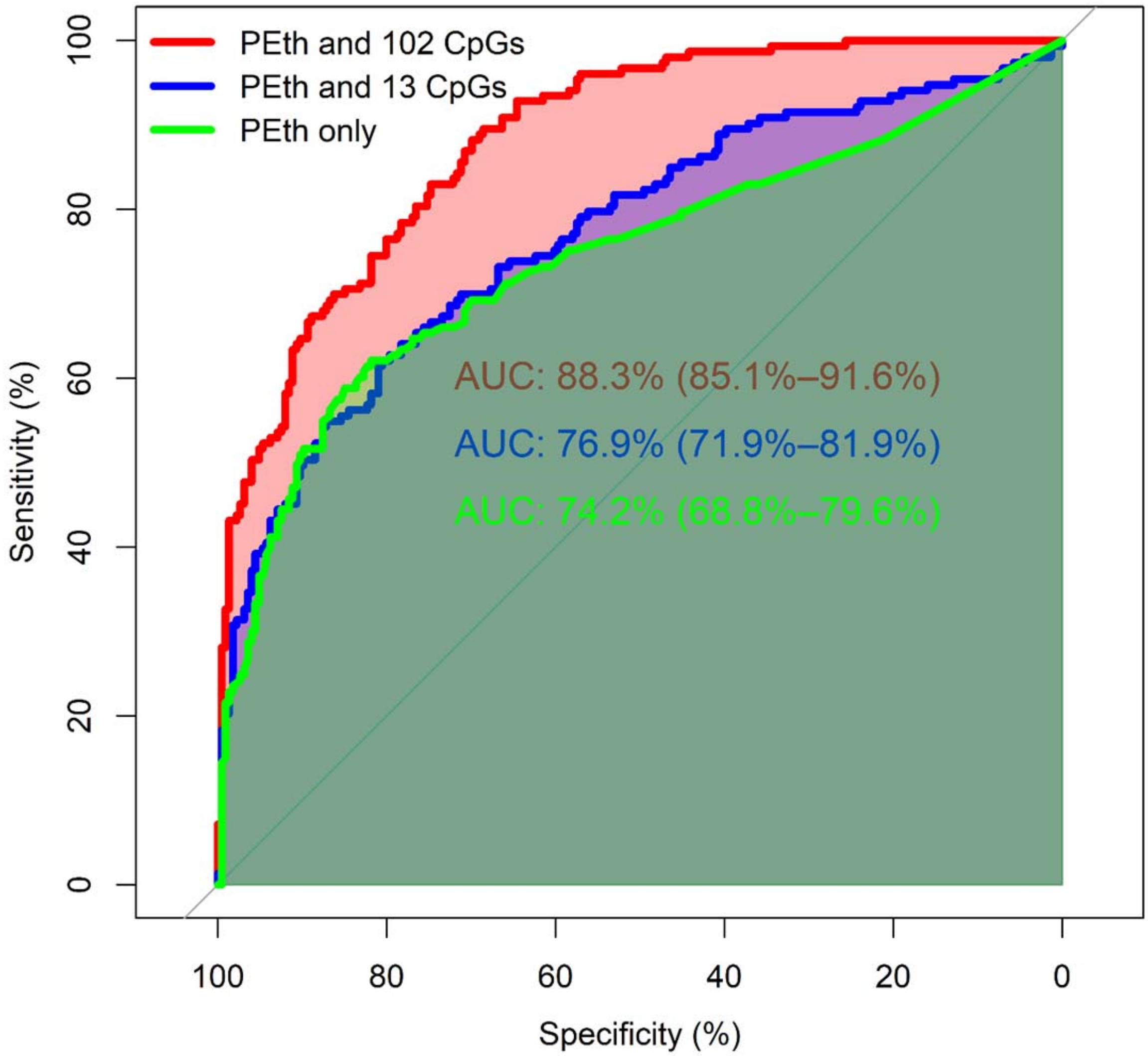

**Figure S4.** Receiver Operating Characteristic (ROC) curve for predicting Hazardous Alcohol Drinking (HAD). ROC curve for predicting HAD by Phosphatidylethanol (PEth) alone, PEth with 13 CpGs (Bonferroni corrected p-value less than 5.00E-02), and PEth with 102 CpGs (false discovery rate (FDR)-corrected p-value less than 5.00E-02) for samples in Cohort 1.

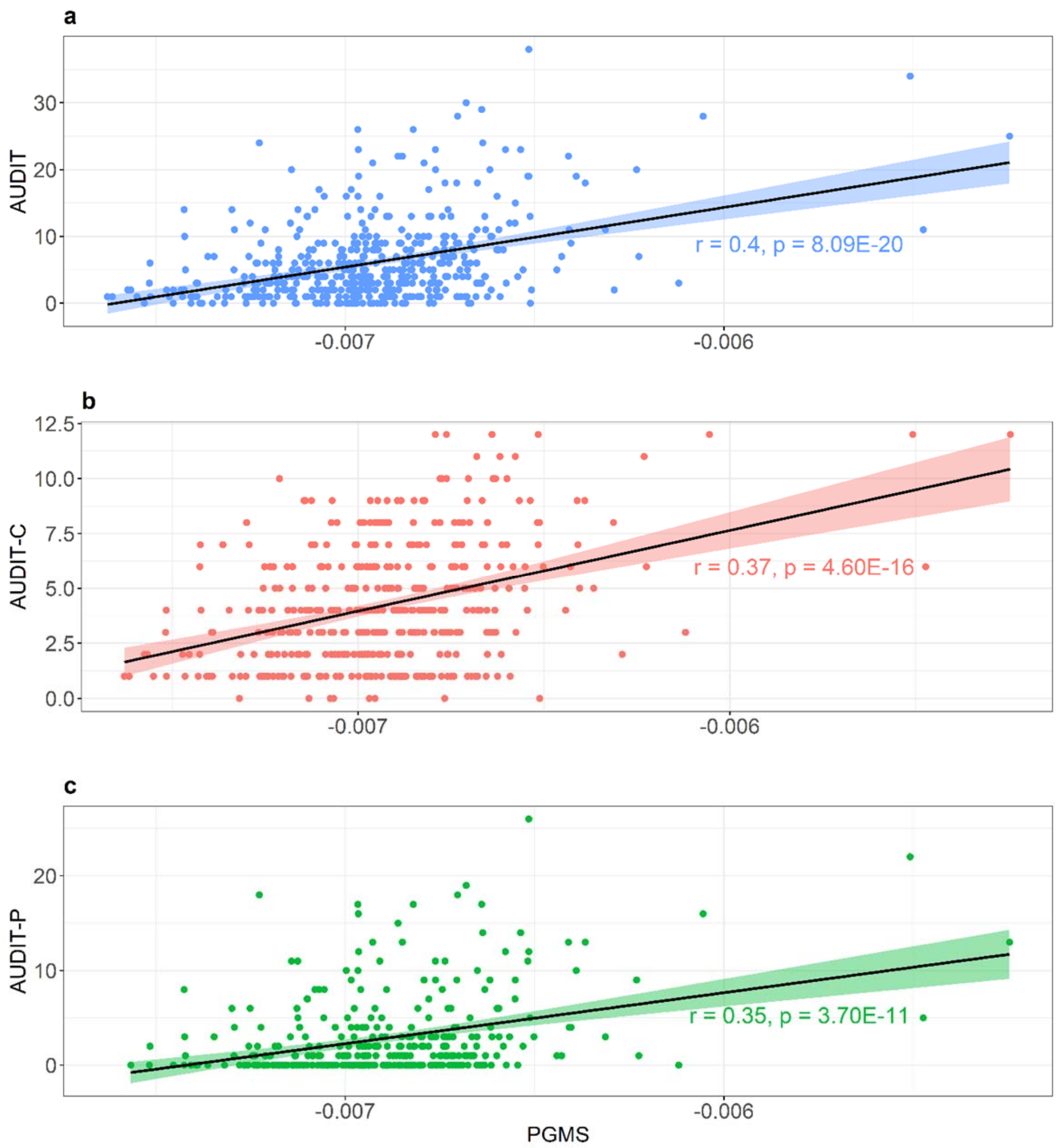

**Figure S5**. Scatterplots for alcohol-related phenotype vs. PolyGenic Methylation Score (PGMS) constructed by 102 Phosphatidylethanol (PEth)-related CpGs. **a.** Scatterplots of Alcohol Use Disorders Identification Test (AUDIT) score vs. PGMS. **b.** Scatterplots of Alcohol Use Disorders Identification Test-Consumption items (AUDIT-C) vs. PGMS. **c.**

Scatterplots of Alcohol Use Disorders Identification Test-Problem items (AUDIT-P) score vs. PGMS.

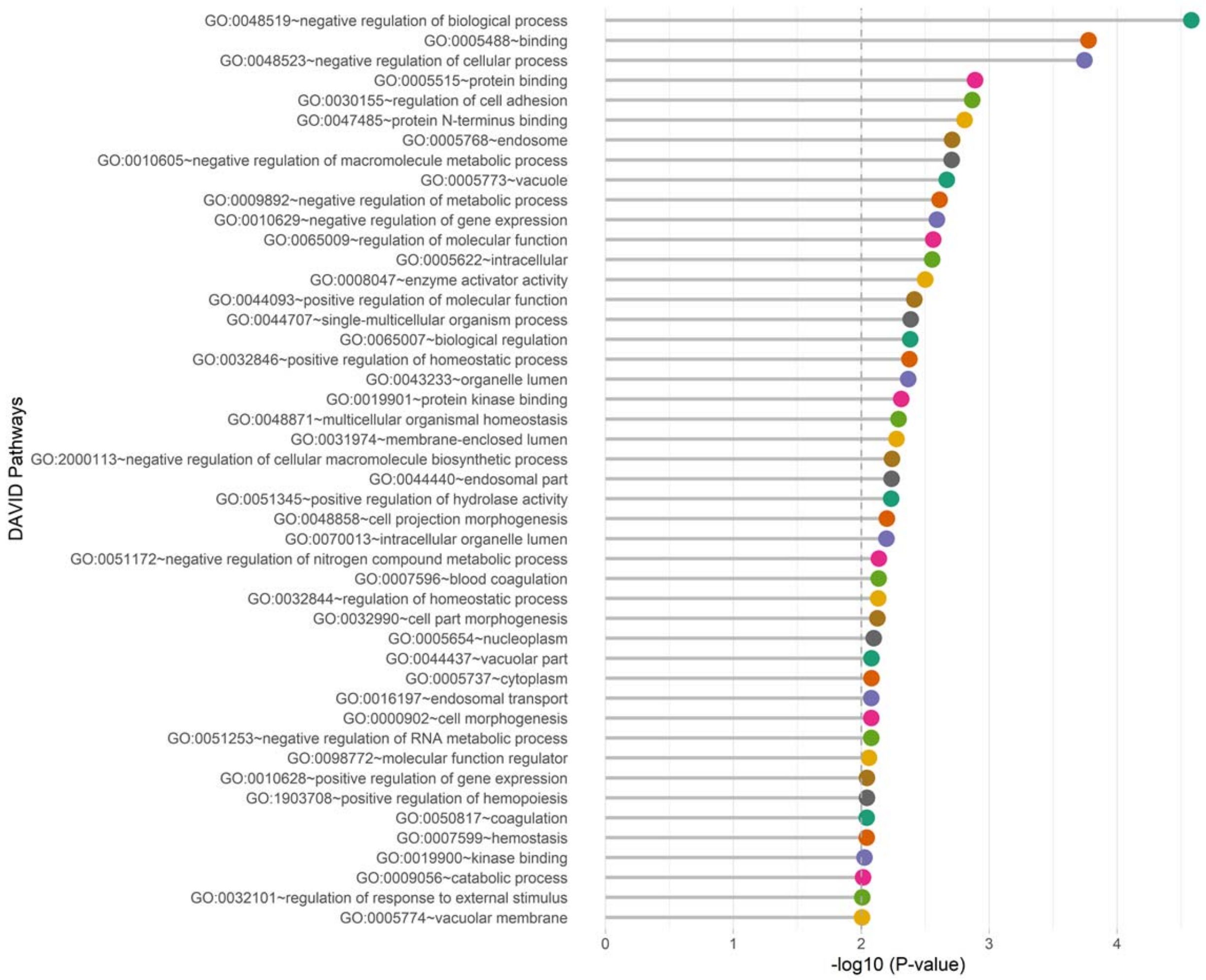

**Figure S6.** Database for Annotation, Visualization and Integrated Discovery (DAVID) pathway analysis for the 130 CpGs selected by elastic net regularization (ENR).
